# Tau4RD fibril polymorphism is imprinted during early aggregation

**DOI:** 10.1101/2025.10.12.681486

**Authors:** Ellie I. James, Mason Saunders, Kelly K. Lee, Miklos Guttman, Abhinav Nath

## Abstract

Microtubule-associated protein tau forms characteristic fibrillar species in many neurodegenerative diseases. Neurofibrillary tangles, tau deposits observed in Alzheimer’s disease (AD), contain a mixture of amyloid-type polymorphic fibrils called paired helical filaments (PHFs) and straight filaments. The formation of heterogenous fibril populations is observed in other diseases and when tau aggregation is induced *in vitro* with polyanionic species. This suggests that tau’s structural transition from a conformational ensemble to various amyloid morphologies is a controlled and, therefore, controllable process. Despite many years of work toward describing aggregation intermediates that could address open questions such as whether fibril polymorphism is imprinted at the start of aggregation or arises due to conformational conversions, our understanding of amyloid structure remains predominantly based on observations of mature fibrils. It is unclear whether these processes are mutually exclusive and to what extent we can bias intermediate conformations toward less toxic states. Here to address the challenge of studying aggregation intermediates and tau’s structural conversion, we apply pulsed hydrogen-deuterium exchange with mass spectrometry (pulsed HDX-MS), which revealed differences in the subpopulations formed by tau4RD (a truncated tau construct) within seconds of initiating aggregation with polyphosphate and within hours of heparin-induction. This work begins to address the gap in knowledge regarding whether amyloid polymorphism is directly imprinted during nucleation or results from structural rearrangement during later stages of aggregation.

## Introduction

Tau and other intrinsically disordered proteins (IDPs) that lack a well-ordered overall conformation are prone to pathological misfolding and aggregation due to their conformational plasticity. The formation and deposition of fibrillar protein aggregates are hallmarks of many neurodegenerative diseases, including tauopathies like Alzheimer’s Disease (AD) and chronic traumatic encephalopathy.^1,2^ In these diseases, microtubule-associated protein tau’s structural role in stabilizing neuronal axons is disrupted and tau begins to self-associate into amyloid-type fibrils with canonical cross-beta sheet morphology. Many of the more than 26 tauopathies described to date exhibit distinct fibrillar aggregates in *ex vivo* samples.^3,4^ Recent studies suggest that the pattern of post-translational modifications (PTMs) on tau is reflective of the tauopathy causing fibrillization.^5,6^ While PTMs are commonly associated with pathological aggregation, the sequence of events leading to PTM enrichment in tauopathies is unclear.^2,7,8^

Amyloidogenic aggregation can be modeled as a defined pathway in which a functional monomer undergoes an activating event that results in an aggregation-competent nucleus to which additional monomers add. This generates oligomers, which further develop into protofibrils and mature fibrils. Inducing tau aggregation *in vitro* often requires phosphorylation or the addition of polyanionic species such as RNA, polyphosphate, or heparin to overcome the positive electrostatic charge spread across the MTBR.^9–11^ Changes in the tau conformational ensemble caused by aggregation initiation are first evident in a short hexapeptide, ^306^VQIVYK^311^, known as PHF6 which has been identified as a nucleation driver. PHF6 is present in all tau isoforms and has been found both necessary and sufficient for tau’s aggregation.^12–14^

Although nucleation begins in the same handful of residues, distinct fibril morphologies result from tau’s incubation with various aggregation inducers.^11,15,16^ In the case of heparin, three fibril polymorphs develop within a single reaction.^16^ Furthermore, aggregation *in vitro* is quite sensitive to changes in parameters such as reaction temperature, buffer composition, and agitation, which results in a broad range of aggregate morphologies even in the presence of a single polyanion.^17,18^ Taken together, these observations suggest that there are points of control along tau’s aggregation pathway that bias the amyloid conformation toward one morphology versus another. This remains an important open question in amyloid research, namely whether amyloid conformations are imprinted at the onset of aggregation or whether differences derive from conformational conversion further along the aggregation pathway.

We have limited insight into the evolution of structure during IDP aggregation, a fact that potentially hampers the development of diagnostics and therapeutics targeted at early intervention. Because oligomers are thought to be toxic in amyloidogenic diseases,^19–24^ it would be of great benefit to therapeutically intervene at the monomeric or intermediate stage and thereby prevent additional neuronal damage. Despite extensive knowledge of fibril morphology due to advances in structural techniques such as cryo-EM,^16,25–27^ this knowledge is not transferable to the dynamic and transient species present during aggregation that elude characterization. Recently, Lövestam *et al*. have pushed the limit by characterizing tau amyloid intermediates by cryo-EM, although their chosen approach provided limited kinetic information.^28^ Pulsed HDX-MS is a solution state technique that is size-agnostic and can inform on the profile of structural order throughout a protein even as it populates transient and disordered species.^29,30^ It can be used to simultaneously identify subpopulations within a reaction (e.g., monomer, oligomer A, oligomer B) and deconvolve aggregation kinetics, facilitating meaningful comparisons about the temporal and structural changes that occur during protein aggregation.^31,32^ Here, we apply pulsed HDX-MS to track the propagation of heparin- or polyphosphate-induced structural differences along tau’s aggregation pathway and find that distinct fibril morphologies appear to be biased during nucleation rather than occurring through a conformational conversion mechanism during mid-to-late aggregation.

## Results

### Generation of conformationally distinct recombinant tau4RD fibrils

It has been shown that inducing tau aggregation with different polyanions generates conformationally distinct fibrillar species.^9,11,27^ Indeed, in our hands, fibrils generated by the addition of heparin or polyphosphate to recombinant tau4RD are morphologically distinct by negative stain EM (nsEM). Differences in branching, helicity, and fibril diameter were observed (Fig. 1A/B, SI Fig. 1). Other factors that could influence fibril morphology, such as temperature, buffer composition, and mechanical agitation, were held constant in these experiments. Limited proteolysis MS/MS experiments supported the inference that tauR4D adopts different conformations since qualitative differences in fibril digestion patterns between monomer, heparin-induced fibrils, and polyphosphate-induced fibrils were clearly evident (SI Fig. 1). We do not interpret the limited proteolysis MS/MS data beyond the qualitative statement that the cleavage products (defined only by c-terminal residue) are different between heparin- and polyphosphate-induced fibrils. While further analysis could reveal additional insight into the amyloid morphologies present in each sample, we do not seek to characterize the fibrils, only their intermediates.

**Figure 1:**
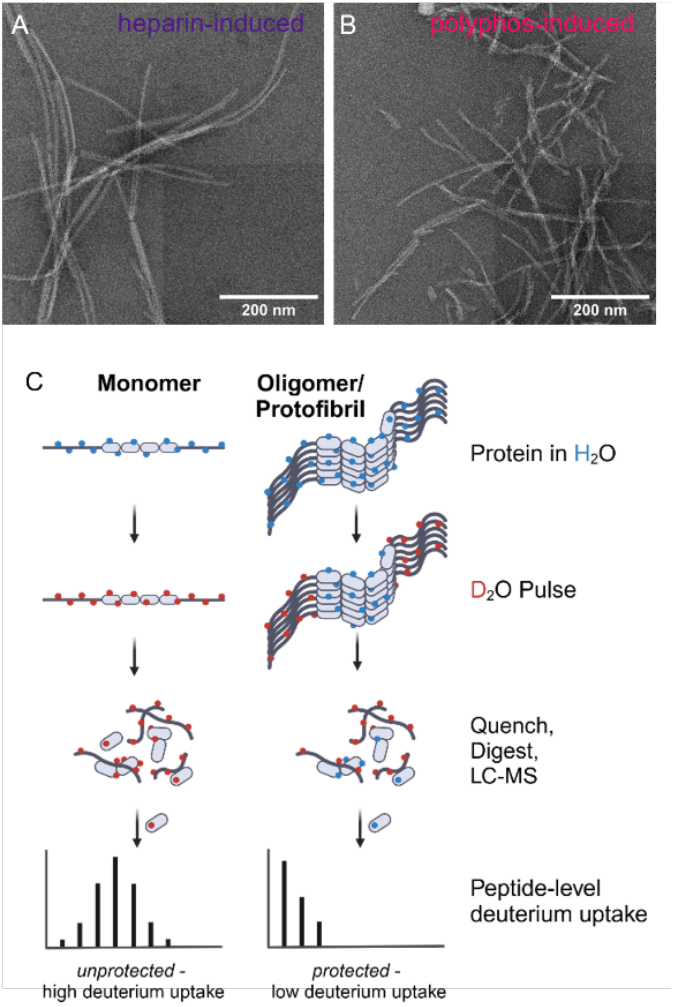
Heparin- and polyphosphate-induced tau4RD aggregation result in different fibril morphologies. A/B) nsEM of representative heparin- and polyphosphate-induced tau4RD fibrils highlighting differences in branching and helicity. Scale bar = 200 nm. C) Cartoon of pulsed HDX experiment showing how differences in protection arise during aggregation.

### Characterization of divergent aggregation pathways

An open question in the amyloid field is whether amyloid structure is imprinted upon aggregation initiation or whether amyloid differentiation occurs through conformational conversion during later stages of the aggregation process. To monitor time-dependent structural changes in tau4RD structure in the presence of polyphosphate or heparin, we employed pulsed HDX-MS. Figure 1C shows a cartoon of a pulsed HDX-MS experiment in which the fibril core is tightly packed and hydrogen bonded, which prevents residues in this region from undergoing HDX. In comparison, the monomer is solvent-exposed and unstructured, making it highly susceptible to HDX. By applying a 3-sec deuterium pulse and observing the peptide-level deuterium uptake, we can discern which regions of tau are monomer-like vs. fibril-like at various points during tau4RD aggregation. Despite attempting to dissociate tau4RD amyloid fibrils with very harsh HDX quench conditions (7 M guanidine HCl), the fibril populations could not be effectively disassembled and were thus not amenable to digestion. Therefore, the populations observed omit the contribution of any fibrils but accurately measure the monomeric and protofibrillar subspecies.

Tau4RD exhibits a decrease in deuterium incorporation during both heparin-induced and polyphosphate-induced aggregation that correlates with amyloid formation monitored by increased ThT fluorescence (Fig. 2A, inset). Tau4RD and inducer concentrations were identical for ThT fluorescence assays and HDX assays, which were performed side-by-side on the same 96-well plate. The only difference in composition between the two assays was the inclusion of 10 μL of ThT solution versus 10 μL of buffer. Without heparin or polyphosphate, there was no decrease in deuterium uptake, indicating that the changes in tau4RD’s deuteration profile are caused by the addition of aggregation inducer (Fig. 2A, dashed grey lines). Protection from HDX and changes in the tau4RD ensemble caused by heparin-induced aggregation appear by 240 min of aggregation (Fig. 2A, inset); at this time, protection appears between residues 284-323 (Fig. 2A, 3D, SI Figs. 2/4). Polyphosphate-induced tau4RD aggregation occurs much more quickly than heparin-induced tau4RD aggregation and lacks a distinct lag phase under our experimental conditions (Fig. 2A, inset). However, the resulting structures are clearly fibrillar (Fig. 1B), and the process is irreversible, indicating that we are observing *bona fide* fibril formation rather than amorphous aggregation or liquid-liquid phase separation. By 15 sec after aggregation initiation (early elongation), pulsed HDX-MS detects protection in peptides ^260^IGS…KKL^282, 291^CGS…VYK^311^, and ^308^IVYKPVDLSK^317^ (Fig. 2A). While Fig. 2A, an alternate visualization of an HDX uptake heatmap, provides a per-residue (averaged) overview of the kinetic and structural differences between heparin- and polyphosphate-induced fibril formation, it obscures key differences in deuteration levels that result from distinct populations in the samples. Figs. 2B/C depict the final pulsed HDX time point for each condition. Protection from HDX (less deuteration) is more widespread throughout the tau4RD sequence (x-axis) during polyphosphate-induced than during heparin-induced aggregation, particularly near the termini. Note the absence of low-exchanging subpopulations in the heparin condition near residues 240-257 and 350-372 (Fig. 2B, SI Fig. 13/14). In both cases, the polyphosphate fibril core extends further than is apparent in Fig. 2A.

**Figure 2:**
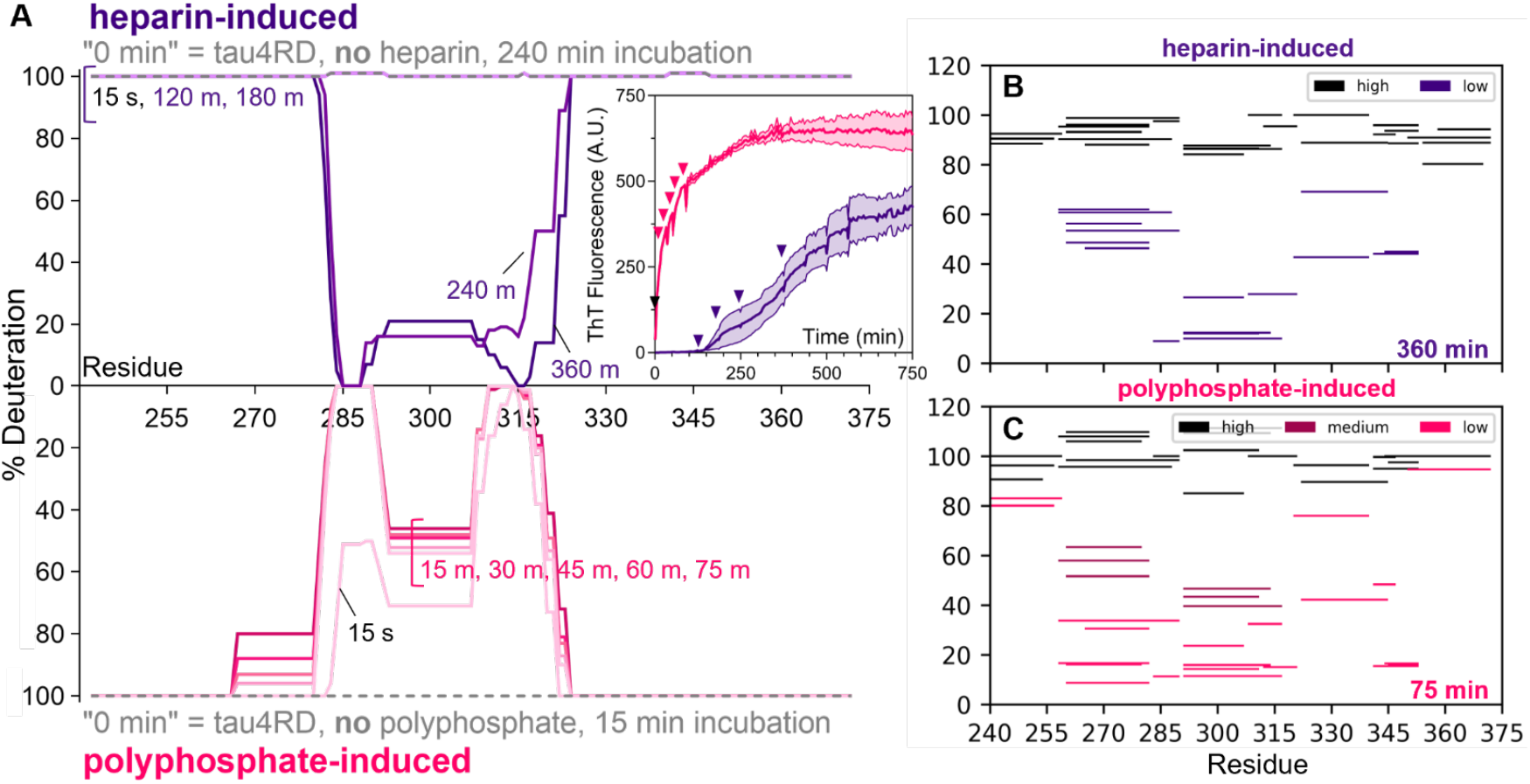
Pulsed HDX-MS detects conformational changes in tau4RD caused by amyloid aggregation. A) Deuteration level per residue of tau4RD following a 3-sec 70% D2O pulse at various aggregation durations. Darker colors indicate longer aggregation. Without inducer (grey dotted lines), tau4RD becomes completely deuterated with a 3-sec D2O pulse. Deuteration levels decrease across the fibril core as aggregation time increases in both the heparin- and polyphosphate-induced conditions. This data primarily reflects the protected (aggregated) population and includes small contributions from the unprotected (monomeric) population. Full peptide coverage information and traditional heat maps can be found in SI Figs. 2-3. Inset: ThT fluorescence monitoring of tau4RD (5 μM) aggregated with 5 μM of 3 kDa heparin or 10 μg/mL medium chain polyphosphate. N = 4, mean ± SEM. Pulsed-HDX sampling times are indicated with colored arrows. B) Woods plot showing deconvoluted % deuteration by peptide for the high-exchanging (monomer-like) and low-exchanging (aggregated) heparin-induced populations at t = 360 m. C) Woods plot showing deconvoluted % deuteration by peptide for the high-exchanging (monomer-like), medium-exchanging (aggregated), and low-exchanging (aggregated) populations at t = 75 m.

One advantage of pulsed HDX-MS is its ability to differentiate the number of conformational subpopulations that exist simultaneously within a sample by quantifying differences in their level of protection. The isotopic distributions in the spectra can reveal the existence of more than the two conformational subpopulations expected when observing a transition like aggregate forming from monomer. HXExpress v3^33^ was used to fit one or more binomial distributions to the mass envelopes of 26 peptides common to heparin- and polyphosphate-induced tau4RD fibrils. Multimodal tau4RD spectra occur throughout the heparin-induced fibril core, including in peptides ^260^IGS…KKL^282^ and ^291^CGS…VYK^311^ (Fig. 3A). These multimodal distributions can be fit with two binomials that represent monomer-like (high-exchanging, unprotected) and fibril-like (low-exchanging, protected) states in peptides across the tau4RD sequence. This single protected species accounts for ∼ 33% of the population 360 min after initiation with heparin (Fig. 3A).

**Figure 3:**
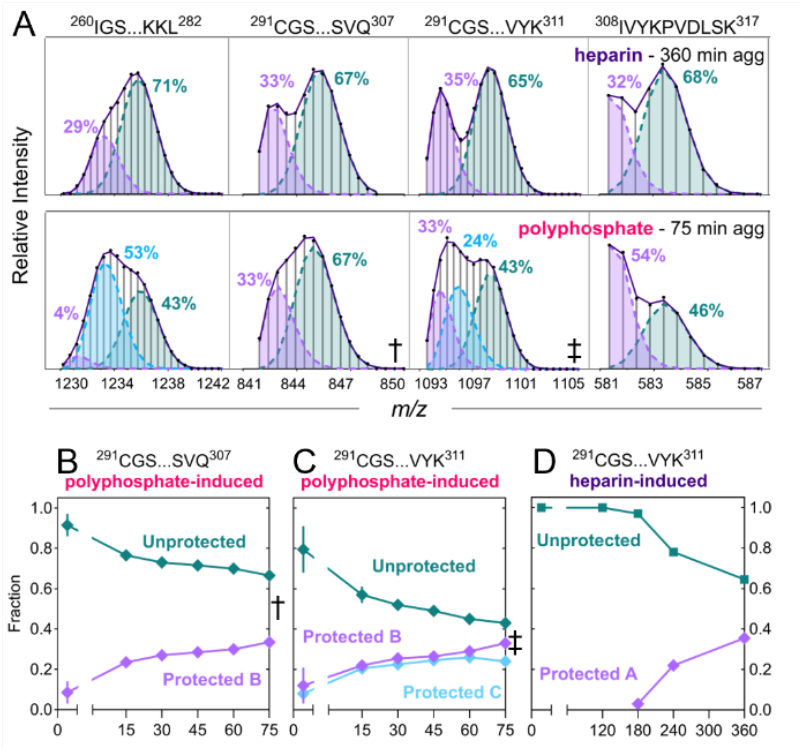
Multimodal deconvolution highlights structural and population differences between heparin- and polyphosphate-induced tau4RD fibril populations. A) Pulsed HDX mass envelopes for five peptides observed in heparin-(top) and polyphosphate-(bottom) induced tau4RD fibrils. In the presence of polyphosphate, peptides ^260^IGS…KKL^282^ and ^291^CGS…VYK^311^ (double dagger) exhibit trimodal character, indicating the existence of at least two stable protected species, while the same peptides display a single protected population in the presence of heparin. The unprotected population decreases over time with a concurrent increase in the protected population(s) under either condition. The F-test p-value for multiple binomial fits is less than 0.05 for all mass envelopes shown. B) Population distribution throughout polyphosphate-induced aggregation for peptide ^291^CGS…SVQ^307^ (dagger) which has been fit to two binomials. C) Population distribution for redundant peptide ^291^CGS…VYK^311^ (double dagger) reveals that a second protected population can be localized to residues near ^308^IVYK^311^ during polyphosphate-induced aggregation. For polyphosphate-induced aggregation, protected B and protected C maintain stable relative population fractions to each other throughout aggregation, indicating that they are two distinct species. The dagger and double dagger symbols mark the time point shown in A. D) In contrast, peptide ^291^CGS…VYK^311^ exhibits a single protected population throughout heparin-induced aggregation. N = 2, mean ± range, except for polyphosphate ^291^CGS…VYK^311^ 75-min (n = 1).

In contrast to our observations of peptides ^260^IGS…KKL^282^ and ^291^CGS…VYK^311^ in heparin-induced tau4RD aggregation, the mass envelope of these peptides cannot be adequately explained by two binomials in the polyphosphate aggregation condition. Applying a multimodal (three binomials) fit to the mass envelope of these peptides results in a statistically justified fit (p-value = 4.0 x 10^-5^, Fig. 3A, SI Fig. 6), indicating the presence of at least two distinct protected populations (i.e., fibril morphs). By comparing the multimodal analyses of peptide ^291^CGS…SVQ^307^ and redundant peptide ^291^CGS…SVQIVYK^311^, we can confidently say that residues ^308^IVYK^311^ in the PHF6 aggregation-driving region are distinct in the two fibril forms (Fig. 3A). Residues beyond the four in this segment may also be distinct in the fibrils but lack of overlapping peptides precludes determining the exact stretch of relevant residues. Further, by comparing the subpopulation distribution between these redundant peptides throughout aggregation, we can determine that at 60 min of aggregation, morph B accounts for 29 ± 0% of the tau4RD population while a second morph accounts for 26 ± 1% of the population and begins in the PHF6 region (Fig. 3B/C). Strikingly, the relative population fractions of these two protected states remain constant throughout aggregation, suggesting that each subpopulation represents a stable protofibril conformation instead of an intermediate species converting into the final conformation (Fig. 3C). The medium- and low-exchanging species seen in Fig. 2C represent separate polyphosphate-induced tau4RD fibril morphs that account for a total of approx. 57% of the population at the final time point.

### Effect of a potent aggregation inhibitor

We have recently demonstrated that tryptanthrin and several synthetic analogs are potent inhibitors of tau4RD aggregation that act primarily by inhibiting tau4RD nucleation.^34^ Here, we investigated whether inhibiting nucleation with tryptanthrin changes the resulting fibril morphologies for either heparin- or polyphosphate-induced tau4RD aggregation. As previously described, the substoichiometric (1:2) addition of tryptanthrin to heparin-induced tau4RD aggregation delayed the onset (longer lag phase) and strongly decreased the rate (decreased transition steepness) and extent (decreased plateau fluorescence) of tau4RD aggregation (Fig. 4A, inset). However, pulsed HDX-MS comparison reveals that tryptanthrin does not cause conformational changes in the core of fibrils formed in the presence of heparin, although peptides at the C-terminal edge of the core shift from weakly multimodal to unimodal (e.g., Fig. 4B/C peptide 341-353) due to the overall reduction in aggregation, resulting in these low abundance populations falling below the limit of detection.

**Figure 4:**
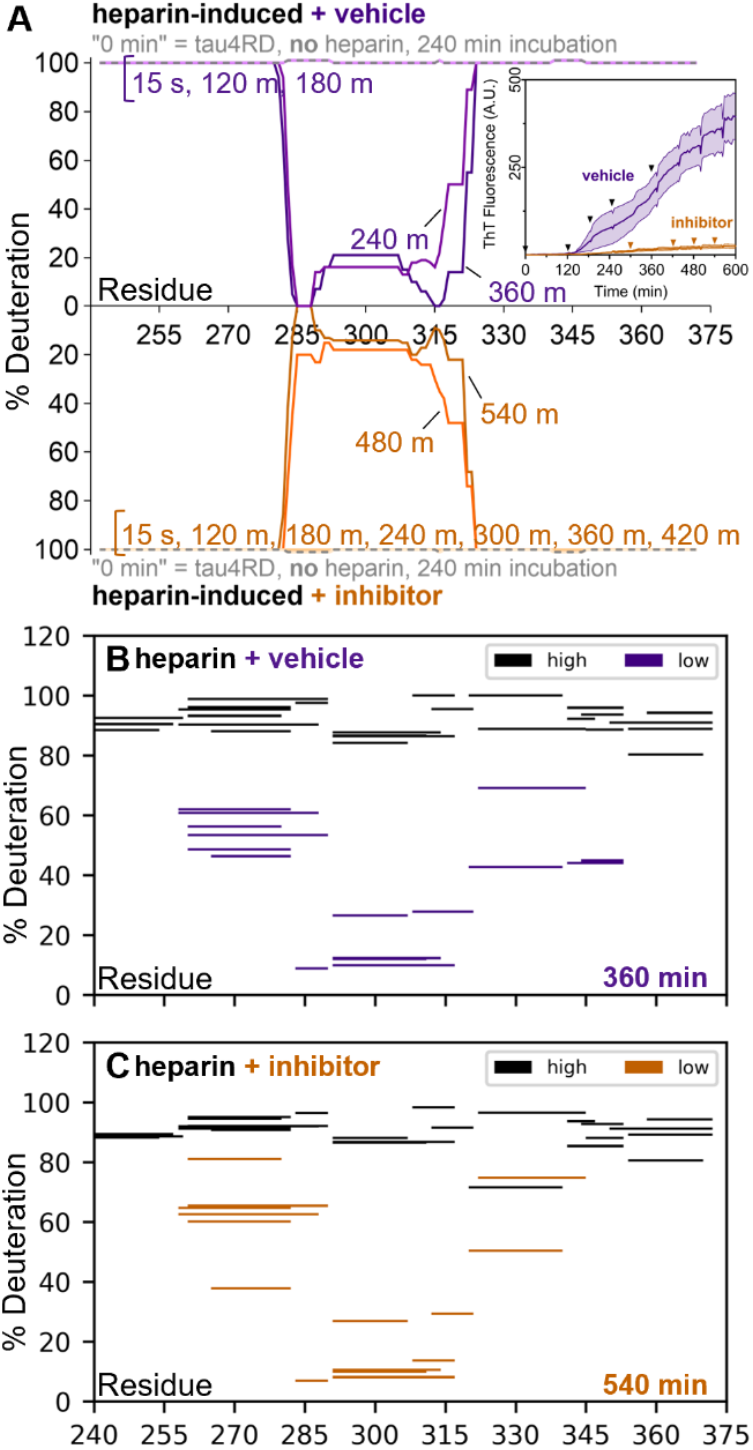
Inhibitor TK23 potently reduces heparin-induced aggregation kinetics but does not change resulting amyloid conformation. A) Deuteration level per residue of tau4RD following a 3-sec 70% D2O pulse at various aggregation durations in the absence or presence of inhibitor TK23. Darker colors indicate longer aggregation. Shared HDX sampling times are indicated in the inset ThT traces with black arrows; condition-specific sampling times are indicated with colored arrows. B/C) Woods plots showing deconvoluted deuterium uptake by peptide for the high-exchanging and low-exchanging (aggregated) populations when incubated with vehicle control (B) or inhibitor (C). Despite the marked decrease in the rate and extent of heparin-induced tau4RD aggregation in the presence of inhibitor TK23 (inset, steepness of curve and decrease in plateau fluorescence), the region protected from HDX remains largely similar to that observed in the absence of inhibitor, except for peptides bordering the fibril core (e.g., 344-353).

Incubation with substoichiometric tryptanthrin (1:2) modestly decreases the extent of polyphosphate-induced tau4RD aggregation (Fig. 5B, inset) but does not result in changes to the fibril core observed by pulsed HDX-MS (Fig. 5B/C). Like the effect observed on the C-terminal side of the heparin fibril core, peptides with low abundance multimodal character (e.g. peptide 350-372, SI Fig. 13) become unimodal after incubation with tryptanthrin. This likely results from the suppression of aggregation rather than a change in the polyphosphate-induced fibril conformations. Moreover, the relative population distribution of morph B and morph C throughout polyphosphate-induced aggregation with tryptanthrin is unchanged (SI Fig. 10), providing additional evidence that the fibril morphs observed during polyphosphate-induced aggregation are unchanged by tryptanthrin and are conformationally distinct from heparin-induced fibrils. Further, it is clear from this data that tryptanthrin does not preferentially inhibit one polyphosphate-induced nucleus compared to the other, which would result in a skewed population distribution compared to the vehicle-controlled state.

**Figure 5:**
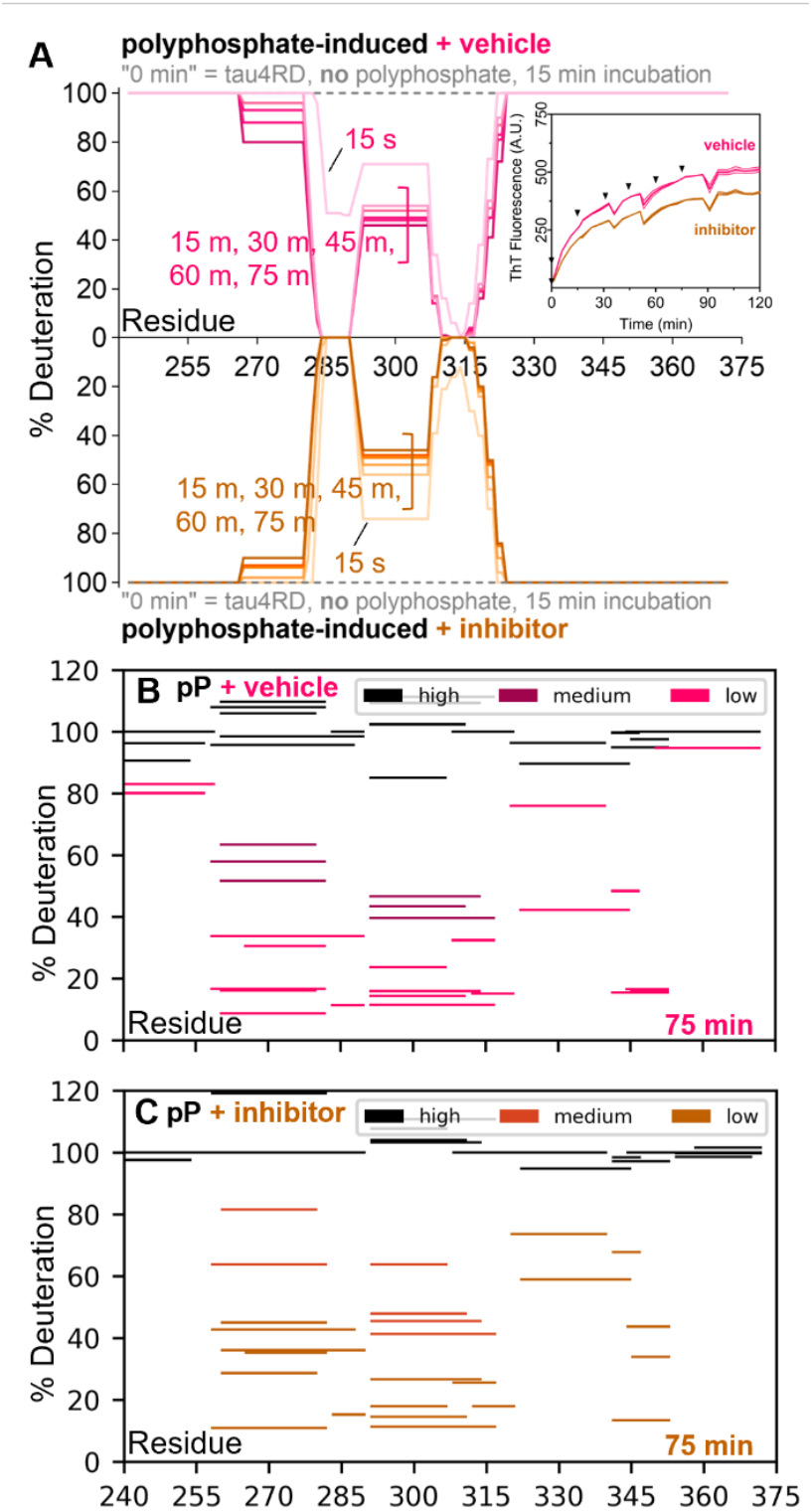
Inhibitor TK23 moderately decreases polyphosphate-induced aggregation but does not change resulting amyloid conformations. A) Deuteration level per residue of tau4RD following a 3-sec 70% D2O pulse at various aggregation durations in the absence or presence of inhibitor TK23. Darker colors indicate longer aggregation. Shared HDX sampling times are indicated in the inset ThT traces with black arrows. Inhibitor TK23 mildly decreases the final extent of polyphosphate-induced tau4RD aggregation (inset, decrease in plateau fluorescence), but has little impact on the rate of aggregation. E/F) Woods plots comparing tau4RD deuterium uptake of the high-(monomer-like), medium-(aggregated), and low-exchanging (aggregated) after aggregation with vehicle (E) or inhibitor (F). The region protected from HDX remains largely unchanged in the presence of TK23, although peptides bordering the fibril core are not detectably multimodal in the presence of inhibitor (e.g., peptide 240-257).

**Figure 6:**
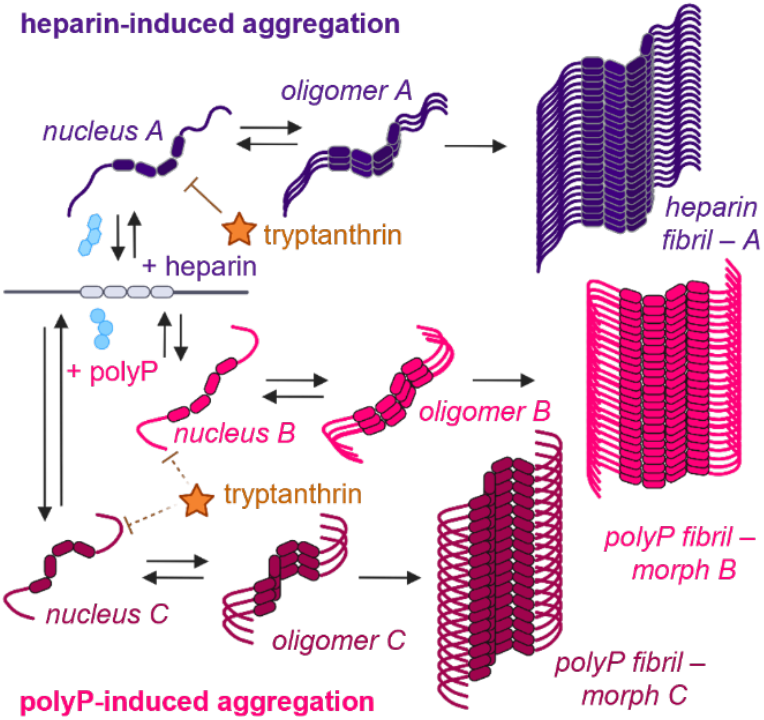
Nucleation proposed as the divergence point of both inter- and intra-inducer tau4RD amyloid polymorphism. Detection of multiple protected populations 15 sec after tau4RD aggregation induction with polyphosphate indicates that fibril polymorphism is imprinted extremely early during aggregation. Tryptanthrin, a potent tau4RD nucleation inhibitor, does not alter the resulting fibril core in heparin- or polyphosphate-induced aggregation. Polymorphic aggregation with stable population fractions is observed when tryptanthrin is incubated with polyphosphate and tau4RD.

## Discussion

Although amyloid aggregation is a complex process, it can be simplified with the application of nomenclature that describes relevant steps in the aggregation process. For example, “nucleation” is the transition that a monomer undergoes to make it aggregation-prone, and “elongation” describes the process by which additional monomers add to existing fibril ends. The microscopic processes involved in amyloid formation (e.g., primary nucleation, fibril fragmentation) are characteristic of an individual protein’s self-assembly landscape and can change depending on the environmental conditions.^17,18,27,35–39^ For example, heparin- and polyphosphate-induced tau4RD aggregation exhibit strikingly different kinetics. Nucleation of tau4RD in the presence of heparin occurs much more slowly than in the presence of polyphosphate, made apparent by the approximately 150-minute lag phase observed in heparin-induced aggregation and the absence of a lag phase during polyphosphate-induced aggregation (Fig. 2 inset). The rate-limiting step of heparin-induced aggregation is nucleation, while that of polyphosphate-induced aggregation is fibril elongation. Although these differences are reflected in the shape of the aggregation curve, they are also represented by the total duration of aggregation. Tau4RD aggregation takes more than twice as long to plateau when initiated with heparin versus polyphosphate, and the shape of the curves indicates a much higher fibril elongation rate (steeper transition slope) observed during polyphosphate-induced aggregation. Although polyphosphate and heparin are both highly negatively charged and have similar charge density per unit length (-0.31 e/Å vs. -0.27 e/ Å), they differ in the chemistry and arrangement of their negative charge (phosphate vs. sulfates/carboxyl).^18^ This results in distinct aggregation kinetics as well as the formation of distinct amyloid species when used to initiate tau4RD aggregation *in vitro*. Both nsEM and limited trypsin/lys-C proteolysis of the polyphosphate- and heparin-induced tau4RD fibrils confirmed that differences exist in the final amyloid structures formed between the two conditions (Fig. 1A/B, SI Fig. 1).

Despite the clear picture of endpoint tau fibril morphologies observed by cryo-electron microscopy,^16,25–27^ the transient nature of amyloid intermediates has, until recently, limited their structural characterization and precluded the determination of when amyloid conformations diverge. Lövestam *et al*. applied time-resolved cryo-EM to monitor the amyloid formation of a truncated tau construct consisting of residues 297-391. The authors discovered a species they termed the FIA (first amyloid intermediate), the first intermediate structured enough to resolve with cryo-EM. During further monitoring, the authors observed maturation and conformational conversion of the FIA into polymorphic intermediates, some of which were shared amongst aggregation conditions.^28^ Although this cryo-EM data provides structural insight into the origin of amyloid polymorphism, it does not provide kinetic details about the assembly reactions.

Pulsed HDX-MS addresses this concern because it can track the kinetics and structural conversions of aggregation *in situ*. Another powerful advantage of pulsed HDX-MS in the study of aggregation intermediates is its ability to discern the prevalence of conformational subpopulations.^29,33^ Unlike the data in Fig. 2A, which is based on the averaged deuterium shift approximated to single residues through all observable peptides, the data in Fig. 3A shows the mass envelope for a given peptide and has been analyzed to discern distinct populations.^33^ This format lends itself to the determination of subpopulations such as the protected (purple, lower mass) and unprotected (teal, higher mass) observed in peptides spanning residues 260-353 for heparin and 240-372 for polyphosphate. Protection extending further than residue 323 is not apparent in the centroided, per-residue deuteration levels in Fig. 2B due to the small and poorly resolved population fractions, emphasizing the importance of multimodal fitting to extract the full scope of information contained within the data. The low abundance protected populations observed at the termini are not artifacts; they are never observed in the monomer injections in which no aggregation has occurred. They also do not appear in earlier time points or in blank injections, which would be expected if they were the result of carryover from fibrils sticking to the protease column and slowly disassembling which would cause apparent protected subpopulations in subsequent runs.

Multimodal fitting of peptide ^291^CGS…VYK^311^ revealed the evolution of multiple protected subpopulations during polyphosphate-induced aggregation and a single protected population during heparin-induced aggregation (Fig. 3A). Remarkably, both polyphosphate-specific protected subpopulations were detected 15 seconds after the addition of polyphosphate and increased in population fraction concurrently (Fig. 3C), lending support to our hypothesis that fibril polymorphism is imprinted early in the aggregation process rather than occurring through a conformational conversion process. This comparison highlights the power of pulsed HDX-MS to interrogate the conformational differences between separate aggregation pathways as well as the differences that develop within a single aggregation reaction.

After establishing pulsed HDX-MS as an experimental tool capable of detecting structural differences between aggregation intermediates, we sought structural and mechanistic information about tryptanthrin-mediated aggregation inhibition. To facilitate detectable levels of aggregation, a substoichiometric (1:2) molar ratio of tryptanthrin to tau4RD was chosen. As expected, heparin-induced aggregation was greatly diminished by the inclusion of tryptanthrin, but somewhat surprisingly, the resulting aggregation intermediates and fibril core were the same as in the control condition. Unlike with heparin, tryptanthrin only mildly inhibits polyphosphate-induced tau4RD aggregation (SI Fig. 7). Once again, tryptanthrin was found not to change the core of either polyphosphate-induced fibril morph or the ratio of their populations to each other, supporting that both polyphosphate-induced polymorphs are distinct from that observed during heparin-induced aggregation and neither is preferentially inhibited by tryptanthrin.

While our data is extremely revealing, there remain some limits to its interpretation. First, polyphosphate-induced tau4RD aggregation proceeds so quickly that there are sizeable changes in the subpopulation fractions of, e.g., peptide ^291^CGS…VYK^311^ between within samples collected in the first minute after initiating aggregation. It was not possible to collect duplicate samples in this time regime due to these changes, so a single data point was collected at 15-sec and 30-sec post-initiation. This limitation is alluded to in Fig. 3B/C and can be seen in more detail in SI Fig. 11. Despite this, both protected subpopulations are visible at the earliest time point in the first replicate (SI Fig. 11), indicating that polymorphism has fully evolved by 15-sec after the start of polyphosphate-induced aggregation.

Second, our pulsed HDX experiments lacked a deuterium exposure control (internal exchange reporter). Although 3-sec deuterium pulses were performed with the aid of a metronome,^40^ there are likely small variations in deuterium exposure between samples. We have mitigated this shortcoming by choosing a pulse duration that adequately separates monomeric and structured tau4RD populations and through our analysis of subpopulation prevalence rather than by comparing %D across samples. Although HXExpress v3 can confidently assign peptide spectra as unimodal, bimodal, trimodal, etc., it cannot deconvolute poorly resolved subpopulation fractions. For this reason, we have taken a conservative approach to inter- and intra-inducer comparisons, focusing on the statistically justified differences in the number of subpopulations detected in key regions of the tau4RD sequence.

Finally, mature amyloid fibrils are not amenable to pulsed HDX-MS in our hands because they cannot be solubilized for LC-MS analysis. We see no changes in the relative signal intensities during LC-MS of peptides in the fibril core versus those at the extreme N- and C-termini in monomeric digests, early aggregation digests, or digests performed during the mid-to-late elongation phase, suggesting that fibrils are not disproportionately contributing to peptide signal at the termini. Negating the contribution of fully formed fibrils means that we do not have a fully quantitative picture of the populations present, but we do still get a complete picture of the early intermediates that lead to fibrils. Fortunately, these are the high-value targets that we wanted to structurally characterize.

Here we have presented a comparative structural analysis of heparin- and polyphosphate-induced tau4RD aggregation. We highlight differences in the extent of the tau4RD sequence involved in the fibril core and differences in conformational polymorphism, specifically observed in regions 260-282 and 291-321 in polyphosphate-induced samples. Our results support the conclusion that tau fibril polymorphism is imprinted during nucleation; we see no evidence of conformational conversion through the course of fibril formation and maturation. This, at first glance, appears to contradict the findings of Lövestam *et al*. who found evidence of an intermediate common to two separate tau aggregation reactions and determined that differences in fibril polymorphism arose through the structural maturation of the FIA. None of the protection profiles we observed are consistent with the FIA, although this is unsurprising due to the different systems our two studies used to induce tau’s aggregation. There are many possible explanations for the disparate conclusions we have reached. Namely, their work and ours made use of different tau constructs which could cause differences in aggregation mechanism. The construct we use (tau4RD, residues 244-372) contains PHF6*, a second hexapeptide motif known to drive aggregation; the construct used by Lövestam *et al*. does not contain PHF6* but does comprise all the residues that form the proteolytically-stable core of PHFs observed in *post-mortem* AD patient samples. The differences in polymorphic behavior we observed could simply be intrinsic to these different tau constructs. Alternatively, the timeline of polyphosphate-induced tau4RD aggregation is far removed from that observed in the Lövestam study or in our own heparin-induced aggregation. It could be that *in vitro* polyphosphate-induced aggregation is kinetically controlled and proceeds without the thermodynamically stable FIA, but conditions that promote an aggregation lag phase allow time for the FIA to arise and mature into separate polymorphs. Regardless, these discrepancies clearly indicate that we need to deepen our understanding of the origin of amyloid polymorphism to create a unified framework.

## Materials and Methods

### Protein Expression and Purification

Tau4RD was generated in-house as recently described. In brief, tau4RD was expressed and purified in *E. coli* BL-21 (DE3) from a plasmid gifted by the Rhoades lab at the University of Pennsylvania (Philadelphia, PA). In brief, a three-step purification consisting of two Ni-NTA agarose column steps followed by fractionation with a 25 mL S200 extended gel filtration column. Tau4RD purity was confirmed by precast 12% Bis-Tris SDS-PAGE gels (Invitrogen). Protein was concentrated, aliquoted, and flash-frozen before storage at -80 °C. Protein concentrations were determined using the Pierce BCA Protein Assay Kit (Thermo Fisher Scientific).

### Aggregation Assays

Tau4RD aggregation assays were completed as previously described with the following changes.^34^ Tau4RD and the aggregation inducers were prepared at 2.105x the desired final concentration in a buffer consisting of 20 mM MOPS, 50 mM NaCl, 1 mM TCEP, 1 mM EDTA mixed in optima H2O, pH = 7.44. The buffer was not filtered prior to use. Tau4RD and tryptanthrin (or an equivalent volume of DMSO) aliquots were thawed and diluted to 10.526 μM (tau4RD) and 5.263 μM (tryptanthrin), respectively, in a combined final volume of 460 μL per experimental condition. For each condition, 95 μL of tau4RD sample was plated into each of n = 3 or 4 wells on a black 96-well plate with a multi-channel pipette. Then, 10 μL of 1 mM filtered (0.22 μm) ThT stock was added to each well with a multi-channel pipette (final concentration of 50 μM ThT). Finally, 95 μL of heparin at 10.526 μM or 95 μL of polyphosphate (Kerafast, medium chain) at 21.05 μg/mL was added to each well for a final concentration of 5 μM heparin or 10 μg/mL polyphosphate (approx. 7 μM when average polyphosphate chain length (mode) is 75 units). The plate was then covered with a polypropylene or polyolefin film and placed into a BioTek Synergy HTXplate reader pre-heated to 37 °C. Aggregation was monitored until completion (overnight, ∼17 h unless stated otherwise) by ThT fluorescence (λ_ex_ = 440 nm, λ_em_ = 485 nm). Fluorescence readings were collected every 5 minutes following 1 minute of linear shaking at 540 rpm. All ThT fluorescence assays were performed with non-binding black 96-well flat-bottom plates (Corning). A blank sample was prepared as above but without tau4RD. The blank was used to correct baseline fluorescence.

### Gel Densitometry

Aggregation assays were stopped at the indicated time and wells were pipetted into individual 1.5 mL tubes. The tubes were centrifuged at 21,000*g* for 30 min at 4 °C, after which 15 μL of supernatant was added to 3 μL of 5x SDS loading dye. Concentration standards were prepared at 5 μM, 2.5 μM, and 1.25 μM and included on the plate during the aggregation assay. Samples and standards were boiled for 45 sec, cooled on ice, then 15 uL was added to precast 12% Bis-Tris SDS-PAGE gels. Gels were imaged with a Li-Cor Odyssey CLx gel scanner and band intensity was determined using GelAnalyzer 23.1.1 (available at www.gelanalyzer.com) by Istvan Lazar Jr., PhD and Istvan Lazar Sr., PhD, CSc. N = 2 or 3 and results were averaged (mean).

### Limited Proteolysis

#### Trypsin/Lys-C Digest

Tau4RD was prepared and aggregated as above, except additional aggregation buffer (10 *μ*L) was added to the wells in place of ThT solution. At the end of aggregation, wells were pipetted individually into non-binding 1.5 mL tubes, then centrifuged at 21,000 g for 30 min at 4 *°C*. Fibril pellets were isolated by removing 175 *μ*L of supernatant from each tube. Trypsin/Lys-C Protease Mix (Pierce, MS-Grade) was resuspended at 0.5 mg/mL in PBS buffer pH 7.4 (20 mM phosphate, 150 mM NaCl, 0.02% sodium azide, 1 mM EDTA), aliquoted, and stored at -20 *°C* until use. For tau4RD fibril digest, 0.4 μg of Trypsin/Lys-C mix was added to each tube. Initially, samples were incubated at 37 *°C* for 15, 30, 60, or 120 min. After comparing cleavage products, a 30-minute digestion was chosen for further experiments. Proteolysis was quenched and fibrils were disrupted with the addition of 8 M urea, pH 2.5, and 30-min incubation at 37 *°C*. Samples were cooled to room temperature, used immediately in an SDS-PAGE gel, or flash-frozen and stored at -80 *°C* until LC-MS/MS analysis.

#### LC-MS/MS

Samples were thawed at 5 °C for 4.5 minutes and injected using a custom LEAP robot integrated with an LC-MS system.41 Peptides were trapped on a Waters XSelect CSH C18 trap cartridge column (2.1 x 5 mm 2.5 µm) and resolved over a CSH C18 column (1 x 50 mm 1.7 µm 130Å) using a linear gradient of 5 to 35% B (A: 0.1% FA, 0.025% TFA, 5% ACN; B: ACN with 0.1% FA) over 10 minutes and analyzed on a Thermo Orbitrap Ascend mass spectrometer at a resolution setting of 120,000. A series of washes over the trap column were used between injections to minimize carry-over as described previously. Data-dependent MS/MS acquisition was performed using rapid CID and HCD scans and processed in Byonic (Protein Metrics) with a digestion filter set to K/R and infinite missed cleavages. A score cutoff of 200 was used to identify peptides.

#### Peptide Mapping

Peptides identified from limited proteolysis-LC-MS/MS were filtered, sorted, and compared using an in-house Python script (available upon request). Briefly, each K/R position in tau4RD was assessed for proteolysis by totaling the XIC of peptides with K/R in the C-terminal position. K/R positions with XIC signals less than 1e5 were assigned an XIC value of 0. The XIC for each K/R position (per sample) was then normalized to the average (mean) XIC of all K/R positions within the sample. This is a simplified workflow that neglects considering the starting residue of each cleavage product. If quantitative rather than qualitative cleavage readouts are necessary, we recommend a more in-depth analysis and comparison.

### Pulsed HDX

Pulsed HDX experiments were designed to minimize perturbing aggregation by separating each time point into a different well on the plate. This means that duplicate samples are created from a single well for each time point, but each time point is pulled from a separate well (i.e., a separate aggregation reaction). As such, the extent of aggregation may vary slightly between time points even though aggregation was initiated at the same time.

Aggregation assays were prepared as described above. Samples for pulsed HDX-MS did not contain ThT (replaced with 10 μL of aggregation buffer), but n = 3 or 4 matched wells contained ThT to monitor aggregation progress during the pulsed HDX experiments. At each time point, the corresponding ThT-free well was transferred into a 1.5 mL lo-bind tube (Eppendorf) and held at 37 °C. The 96-well plate was returned to the plate reader for continued monitoring. Then, 20 μL of tau4RD/mixture was diluted into 80 μL of deuterated MOPS buffer (20 mM MOPS, 50 mM NaCl, 1 mM TCEP, 1 mM EDTA, 0.2 nM bradykinin, pH* 7.18, 70% D final) for 3 sec at 21 °C. Timing accuracy was facilitated using an electronic metronome set to 60 beats/min. After 3 sec, the sample was rapidly mixed with an equal volume of ice-cold quench (4 M urea, 0.2 % formic acid, 0.1% TFA) for a final pH of 2.51. Samples were immediately frozen on ethanol/dry ice and stored at -80 °C until LC-MS analysis. Samples were collected in duplicate at each aggregation time point. Fully deuterated samples were prepared by incubating monomeric tau4RD with deuterated buffer at 21 °C for 60 min. Samples were then quenched and frozen as above. Undeuterated samples were prepared alongside fully deuterated samples but with a matched deuterium-free buffer.

### LC-MS for HDX samples

Of note, samples containing polyphosphate were challenging to assay via LC-MS and significant effort went into identifying suitable experimental conditions. Briefly, polyphosphate buildup on the trap and analytical columns prevented peptide binding, resulting in blank or extremely noisy mass spectra. A neutral-pH wash was passed over the trap and analytical columns between each sample injection to remove polyphosphate buildup. No carryover was detected in blanks injected between polyphosphate-or heparin-containing samples.

All HDX samples were subjected to LC-MS on a Thermo Orbitrap Ascend during an uninterrupted three-day period. Samples were thawed at 5 °C for 4.5 minutes and injected using a custom LEAP robot integrated with an LC-MS system.^41^ The protein was first passed over a Nepenthesin II column (2.1 x 30 mm; AffiPro) at 400 µL/min for inline digestion with the protease column held at 20 °C. Peptides were then trapped, eluted, and analyzed as described above. A series of washes over the trap and protease columns was used between injections to minimize carry-over as described previously,^41^ except the fourth wash which was replaced with 98% optima water/2% optima ACN (neutral pH). Data-dependent MS/MS acquisition was performed on an undeuterated sample using rapid CID and HCD scans and processed in Byonic (Protein Metrics) with a score cutoff of 200 to identify peptides. Deuterium incorporation was analyzed using HDExaminer v3 (Trajan Scientific). Peptide spectra were exported from HDExaminer and analyzed with HXExpress v3 for multimodal character.^33^ Multiple binomials were fit to spectra based on the F-test p-value calculated in HXExpress v3. Multimodal spectra were plotted according to the solutions provided by HXExpress v3, regardless of the confidence associated with the subpopulation deconvolution. Only well-resolved or redundant spectra were used to assign population fractions.

### nsEM

Tau4RD was prepared, aggregated, and pelleted as for limited proteolysis. Fibril pellets were isolated by removing 190 uL of supernatant from each tube. Fibrils were resuspended by gently pipetting the remaining 10 μL of buffer until the pellet was dispersed. Negative stain grids were made by applying 3.5 μl of dispersed fibrils to carbon film 400 mesh copper grids (Electron Microscopy Sciences) for one minute, blotting to dryness, washing with distilled water, blotting to dryness, staining using a 2% methylammonium tungstate solution (NanoW™; Nanoprobes) for one minute, and then blotting to dryness. nsEM was performed on a 120 kV Tecnai T12 TEM (FEI). Micrographs were collected at 67,000x magnification with a 1.60 Å/pixel size using the software Leginon^42^ with defocus set to -2.0 μm and a total electron dose of 35.04 e−/Å.

## Acknowledgments

The authors wish to thank Dale Whittington and Natalie Stone for assistance with data collection. This work was funded by National Institutes of Health grants: R01GM127579 (M.G.), T32GM007750 (E.I.J.), S10OD030237 (M.G.), and the National Science Foundation Award: 2304707 (M.G.). The authors gratefully acknowledge support from the Seattle Partnership for Research on Innovative Therapies (to A.N.) and the Sidney Nelson Endowment (to K.K.L.).

## Author contributions

E.I.J, M.G., and A.N. conceived the project. E.I.J performed *in vitro* aggregation assays, limited proteolysis, and pulsed HDX. M.S. generated nsEM data, which was analyzed by M.S. and K.K.L. E.I.J. drafted the manuscript with input from K.K.L., M.G., and A.N.

## Notes

### Competing Interest Statement

The authors have declared no competing interest.

## References

(1) Zhang, Y.; Wu, K. M.; Yang, L.; Dong, Q.; Yu, J. T. Tauopathies: New Perspectives and Challenges. Mol. Neurodegener. 2022, 17 (1). 10.1186/s13024-022-00533-z.

(2) Limorenko, G.; Lashuel, H. A. Revisiting the Grammar of Tau Aggregation and Pathology Formation: How New Insights from Brain Pathology Are Shaping How We Study and Target Tauopathies. Chem. Soc. Rev. 2022, 51 (2), 513–565. 10.1039/D1CS00127B.

(3) Shi, Y.; Zhang, W.; Yang, Y.; Murzin, A. G.; Falcon, B.; Kotecha, A.; van Beers, M.; Tarutani, A.; Kametani, F.; Garringer, H. J.; Vidal, R.; Hallinan, G. I.; Lashley, T.; Saito, Y.; Murayama, S.; Yoshida, M.; Tanaka, H.; Kakita, A.; Ikeuchi, T.; Robinson, A. C.; Mann, D. M. A.; Kovacs, G. G.; Revesz, T.; Ghetti, B.; Hasegawa, M.; Goedert, M.; Scheres, S. H. W. Structure-Based Classification of Tauopathies. Nature 2021, 598 (7880), 359. 10.1038/S41586-021-03911-7.

(4) Kovacs, G. G.; Ghetti, B.; Goedert, M. Classification of Diseases with Accumulation of Tau Protein. Neuropathol. Appl. Neurobiol. 2022, 48 (3), e12792. 10.1111/NAN.12792.

(5) Wesseling, H.; Mair, W.; Kumar, M.; Schlaffner, C. N.; Tang, S.; Beerepoot, P.; Fatou, B.; Guise, A. J.; Cheng, L.; Takeda, S.; Muntel, J.; Rotunno, M. S.; Dujardin, S.; Davies, P.; Kosik, K. S.; Miller, B. L.; Berretta, S.; Hedreen, J. C.; Grinberg, L. T.; Seeley, W. W.; Hyman, B. T.; Steen, H.; Steen, J. A. Tau PTM Profiles Identify Patient Heterogeneity and Stages of Alzheimer’s Disease. Cell 2020, 183 (6), 1699-1713.e13. 10.1016/J.CELL.2020.10.029.

(6) Kyalu Ngoie Zola, N.; Balty, C.; Pyr dit Ruys, S.; Vanparys, A. A. T.; Huyghe, N. D. G.; Herinckx, G.; Johanns, M.; Boyer, E.; Kienlen-Campard, P.; Rider, M. H.; Vertommen, D.; Hanseeuw, B. J. Specific Post-Translational Modifications of Soluble Tau Protein Distinguishes Alzheimer’s Disease and Primary Tauopathies. Nat. Commun. 2023 141 2023, 14 (1), 1–17. 10.1038/S41467-023-39328-1.

(7) Mirbaha, H.; Chen, D.; Mullapudi, V.; Terpack, S. J.; White, C. L.; Joachimiak, L. A.; Diamond, M. I. Seed-Competent Tau Monomer Initiates Pathology in a Tauopathy Mouse Model. J. Biol. Chem. 2022, 298 (8), 102163. 10.1016/J.JBC.2022.102163.

(8) Li, L.; Nguyen, B. A.; Mullapudi, V.; Li, Y.; Saelices, L.; Joachimiak, L. A. Disease-Associated Patterns of Acetylation Stabilize Tau Fibril Formation. Structure 2023, 31 (9), 1025-1037.e4. 10.1016/J.STR.2023.05.020.

(9) Wickramasinghe, S. P.; Lempart, J.; Merens, H. E.; Murphy, J.; Huettemann, P.; Jakob, U.; Rhoades, E. Polyphosphate Initiates Tau Aggregation through Intra- and Intermolecular Scaffolding. Biophys. J. 2019, 117 (4), 717–728. 10.1016/J.BPJ.2019.07.028.

(10) Chakraborty, P.; Rivière, G.; Liu, S.; de Opakua, A. I.; Dervişoğlu, R.; Hebestreit, A.; Andreas, L. B.; Vorberg, I. M.; Zweckstetter, M. Co-Factor-Free Aggregation of Tau into Seeding-Competent RNA-Sequestering Amyloid Fibrils. Nat. Commun. 2021, 12 (1). 10.1038/s41467-021-24362-8.

(11) Abskharon, R.; Sawaya, M. R.; Cao, Q.; Nguyen, B. A.; Boyer, D. R.; Cascio, D.; Eisenberg, D. S. Cryo-EM Structure of RNA-Induced Tau Fibrils Reveals a Small C-Terminal Core That May Nucleate Fibril Formation. Proc. Natl. Acad. Sci. U. S. A. 2022, 119 (15), e2119952119. 10.1073/PNAS.2119952119/SUPPL_FILE/PNAS.2119952119.SAPP.PDF.

(12) Von Bergen, M.; Friedhoff, P.; Biernat, J.; Heberle, J.; Mandelkow, E. M.; Mandelkow, E. Assembly of τ Protein into Alzheimer Paired Helical Filaments Depends on a Local Sequence Motif (306VQIVYK311) Forming β Structure. Proc. Natl. Acad. Sci. U. S. A. 2000, 97 (10), 5129–5134. 10.1073/PNAS.97.10.5129/ASSET/7E919641-17A6-4468-A582-4672B009D09E/ASSETS/GRAPHIC/PQ0904029008.JPEG.

(13) Ganguly, P.; Do, T. D.; Larini, L.; Lapointe, N. E.; Sercel, A. J.; Shade, M. F.; Feinstein, S. C.; Bowers, M. T.; Shea, J. E. Tau Assembly: The Dominant Role of PHF6 (VQIVYK) in Microtubule Binding Region Repeat R3. J. Phys. Chem. B 2015, 119 (13), 4582–4593. 10.1021/ACS.JPCB.5B00175/SUPPL_FILE/JP5B00175_SI_001.PDF.

(14) Smit, F. X.; Luiken, J. A.; Bolhuis, P. G. Primary Fibril Nucleation of Aggregation Prone Tau Fragments PHF6 and PHF6. J. Phys. Chem. B 2017, 121 (15), 3250–3261. 10.1021/ACS.JPCB.6B07045/ASSET/IMAGES/LARGE/JP-2016-070454_0013.JPEG.

(15) Despres, C.; Di, J.; Cantrelle, F. X.; Li, Z.; Huvent, I.; Chambraud, B.; Zhao, J.; Chen, J.; Chen, S.; Lippens, G.; Zhang, F.; Linhardt, R.; Wang, C.; Klärner, F. G.; Schrader, T.; Landrieu, I.; Bitan, G.; Smet-Nocca, C. Major Differences between the Self-Assembly and Seeding Behavior of Heparin-Induced and in Vitro Phosphorylated Tau and Their Modulation by Potential Inhibitors. ACS Chem. Biol. 2019, 14 (6), 1363–1379. 10.1021/ACSCHEMBIO.9B00325/SUPPL_FILE/CB9B00325_SI_001.PDF.

(16) Zhang, W.; Falcon, B.; Murzin, A. G.; Fan, J.; Crowther, R. A.; Goedert, M.; Scheres, S. H. W. Heparin-Induced Tau Filaments Are Polymorphic and Differ from Those in Alzheimer’s and Pick’s Diseases. eLife 2019, 8. 10.7554/ELIFE.43584.

(17) Axell, E.; Hu, J.; Lindberg, M.; Dear, A. J.; Ortigosa-Pascual, L.; Andrzejewska, E. A.; Šneideriene, G.; Thacker, D.; Knowles, T. P. J.; Sparr, E.; Linse, S. The Role of Shear Forces in Primary and Secondary Nucleation of Amyloid Fibrils. Proc. Natl. Acad. Sci. U. S. A. 2024, 121 (25), e2322572121. 10.1073/PNAS.2322572121/SUPPL_FILE/PNAS.2322572121.SAPP.PDF.

(18) Montgomery, K. M.; Carroll, E. C.; Thwin, A. C.; Quddus, A. Y.; Hodges, P.; Southworth, D. R.; Gestwicki, J. E. Chemical Features of Polyanions Modulate Tau Aggregation and Conformational States. J. Am. Chem. Soc. 2023, 145 (7), 3926–3936. 10.1021/jacs.2c08004.

(19) Emin, D.; Zhang, Y. P.; Lobanova, E.; Miller, A.; Li, X.; Xia, Z.; Dakin, H.; Sideris, D. I.; Lam, J. Y. L.; Ranasinghe, R. T.; Kouli, A.; Zhao, Y.; De, S.; Knowles, T. P. J.; Vendruscolo, M.; Ruggeri, F. S.; Aigbirhio, F. I.; Williams-Gray, C. H.; Klenerman, D. Small Soluble α-Synuclein Aggregates Are the Toxic Species in Parkinson’s Disease. Nat. Commun. 2022 131 2022, 13 (1), 1–15. 10.1038/S41467-022-33252-6.

(20) Ward, S. M.; Himmelstein, D. S.; Lancia, J. K.; Binder, L. I. Tau Oligomers and Tau Toxicity in Neurodegenerative Disease. Biochem. Soc. Trans. 2012, 40 (4), 667. 10.1042/BST20120134.

(21) Ondrejcak, T.; Hu, N. W.; Qi, Y.; Klyubin, I.; Corbett, G. T.; Fraser, G.; Perkinton, M. S.; Walsh, D. M.; Billinton, A.; Rowan, M. J. Soluble Tau Aggregates Inhibit Synaptic Long-Term Depression and Amyloid β-Facilitated LTD in Vivo. Neurobiol. Dis. 2019, 127, 582–590. 10.1016/j.nbd.2019.03.022.

(22) Lee, M. C.; Yu, W. C.; Shih, Y. H.; Chen, C. Y.; Guo, Z. H.; Huang, S. J.; Chan, J. C. C.; Chen, Y. R. Zinc Ion Rapidly Induces Toxic, off-Pathway Amyloid-β Oligomers Distinct from Amyloid-β Derived Diffusible Ligands in Alzheimer’s Disease. Sci. Rep. 2018 81 2018, 8 (1), 1–16. 10.1038/S41598-018-23122-X.

(23) Lo Cascio, F.; Puangmalai, N.; Ellsworth, A.; Bucchieri, F.; Pace, A.; Palumbo Piccionello, A.; Kayed, R. Toxic Tau Oligomers Modulated by Novel Curcumin Derivatives. Sci. Rep. 2019 91 2019, 9 (1), 1–16. 10.1038/S41598-019-55419-W.

(24) Almeida, Z. L.; Brito, R. M. M. Structure and Aggregation Mechanisms in Amyloids. Molecules 2020, 25 (5), 1195. 10.3390/MOLECULES25051195.

(25) Fitzpatrick, A. W. P.; Falcon, B.; He, S.; Murzin, A. G.; Murshudov, G.; Garringer, H. J.; Crowther, R. A.; Ghetti, B.; Goedert, M.; Scheres, S. H. W. Cryo-EM Structures of Tau Filaments from Alzheimer’s Disease. Nature 2017, 547 (7662), 185–190. 10.1038/NATURE23002.

(26) Falcon, B.; Zhang, W.; Murzin, A. G.; Murshudov, G.; Garringer, H. J.; Vidal, R.; Crowther, R. A.; Ghetti, B.; Scheres, S. H. W.; Goedert, M. Structures of Filaments from Pick’s Disease Reveal a Novel Tau Protein Fold. Nature 2018, 561 (7721), 137–140. 10.1038/S41586-018-0454-Y.

(27) Lövestam, S.; Koh, F. A.; van Knippenberg, B.; Kotecha, A.; Murzin, A. G.; Goedert, M.; Scheres, S. H. W. Assembly of Recombinant Tau into Filaments Identical to Those of Alzheimer’s Disease and Chronic Traumatic Encephalopathy. eLife 2022, 11. 10.7554/eLife.76494.

(28) Lövestam, S.; Li, D.; Wagstaff, J. L.; Kotecha, A.; Kimanius, D.; McLaughlin, S. H.; Murzin, A. G.; Freund, S. M. V.; Goedert, M.; Scheres, S. H. W. Disease-Specific Tau Filaments Assemble via Polymorphic Intermediates. Nat. 2023 6257993 2023, 625 (7993), 119–125. 10.1038/S41586-023-06788-W.

(29) James, E. I.; Murphree, T. A.; Vorauer, C.; Engen, J. R.; Guttman, M. Advances in Hydrogen/Deuterium Exchange Mass Spectrometry and the Pursuit of Challenging Biological Systems. Chem. Rev. 2022, 122 (8), 7562–7623. 10.1021/ACS.CHEMREV.1C00279.

(30) Kumar, H.; Udgaonkar, J. B. Mechanistic and Structural Origins of the Asymmetric Barrier to Prion-like Cross-Seeding between Tau-3R and Tau-4R. J. Mol. Biol. 2018, 430 (24), 5304–5312. 10.1016/J.JMB.2018.09.010.

(31) Illes-Toth, E.; Rempel, D. L.; Gross, M. L. Pulsed Hydrogen-Deuterium Exchange Illuminates the Aggregation Kinetics of α-Synuclein, the Causative Agent for Parkinson’s Disease. ACS Chem. Neurosci. 2018, 9 (6), 1469– 1476. 10.1021/ACSCHEMNEURO.8B00052/ASSET/IMAGES/MEDIUM/CN-2018-00052K_M006.GIF.

(32) Illes-Toth, E.; Meisl, G.; Rempel, D. L.; Knowles, T. P. J.; Gross, M. L. Pulsed Hydrogen-Deuterium Exchange Reveals Altered Structures and Mechanisms in the Aggregation of Familial Alzheimer’s Disease Mutants. ACS Chem. Neurosci. 2021, 12 (11), 1972–1982. 10.1021/ACSCHEMNEURO.1C00072/ASSET/IMAGES/LARGE/CN1C00072_0005.JPEG.

(33) Tuttle, L. M.; James, E. I.; Georgescauld, F.; Wales, T. E.; Weis, D. D.; Engen, J. R.; Nath, A.; Klevit, R. E.; Guttman, M. Rigorous Analysis of Multimodal HDX-MS Spectra. J. Am. Soc. Mass Spectrom. 2025. 10.1021/JASMS.4C00471/ASSET/IMAGES/LARGE/JS4C00471_0005.JPEG.

(34) James, E. I.; Baggett, D. W.; Chang, E.; Schachter, J.; Nixey, T.; Choi, K.; Guttman, M.; Nath, A. Tryptanthrin Analogs Substoichiometrically Inhibit Seeded and Unseeded Tau4RD Aggregation. eLife 2024, 13. 10.7554/ELIFE.98227.1.

(35) Tanaka, G.; Yamanaka, T.; Furukawa, Y.; Kajimura, N.; Mitsuoka, K.; Nukina, N. Sequence-and Seed-Structure-Dependent Polymorphic Fibrils of Alpha-Synuclein. Biochim. Biophys. Acta BBA - Mol. Basis Dis. 2019, 1865 (6), 1410–1420. 10.1016/J.BBADIS.2019.02.013.

(36) Yoo, J. M.; Lin, Y.; Heo, Y.; Lee, Y. H. Polymorphism in Alpha-Synuclein Oligomers and Its Implications in Toxicity under Disease Conditions. Front. Mol. Biosci. 2022, 9, 959425. 10.3389/FMOLB.2022.959425/BIBTEX.

(37) Meisl, G.; Rajah, L.; Cohen, S. A. I.; Pfammatter, M.; Šarić, A.; Hellstrand, E.; Buell, A. K.; Aguzzi, A.; Linse, S.; Vendruscolo, M.; Dobson, C. M.; Knowles, T. P. J. Scaling Behaviour and Rate-Determining Steps in Filamentous Self-Assembly. Chem. Sci. 2017, 8 (10), 7087–7097. 10.1039/c7sc01965c.

(38) Knowles, T. P. J.; Waudby, C. A.; Devlin, G. L.; Cohen, S. I. A.; Aguzzi, A.; Vendruscolo, M.; Terentjev, E. M.; Welland, M. E.; Dobson, C. M. An Analytical Solution to the Kinetics of Breakable Filament Assembly. Science 2009, 326 (5959), 1533–1537. 10.1126/SCIENCE.1178250.

(39) Cohen, S. I. A.; Vendruscolo, M.; Dobson, C. M.; Knowles, T. P. J. From Macroscopic Measurements to Microscopic Mechanisms of Protein Aggregation. J. Mol. Biol. 2012, 421 (2–3), 160–171. 10.1016/j.jmb.2012.02.031.

(40) Benhaim, M. A.; Mangala Prasad, V.; Garcia, N. K.; Guttman, M.; Lee, K. K. Structural Monitoring of a Transient Intermediate in the Hemagglutinin Fusion Machinery on Influenza Virions. Sci. Adv. 2020, 6 (18). 10.1126/SCIADV.AAZ8822/SUPPL_FILE/AAZ8822_SM.PDF.

(41) Watson, M. J.; Harkewicz, R.; Hodge, E. A.; Vorauer, C.; Palmer, J.; Lee, K. K.; Guttman, M. Simple Platform for Automating Decoupled LC-MS Analysis of Hydrogen/Deuterium Exchange Samples. J. Am. Soc. Mass Spectrom. 2021, 32 (2), 597–600. 10.1021/JASMS.0C00341/SUPPL_FILE/JS0C00341_SI_001.PDF.

(42) Suloway, C.; Pulokas, J.; Fellmann, D.; Cheng, A.; Guerra, F.; Quispe, J.; Stagg, S.; Potter, C. S.; Carragher, B. Automated Molecular Microscopy: The New Leginon System. J. Struct. Biol. 2005, 151 (1), 41–60. 10.1016/j.jsb.2005.03.010.

