## Supplemental Data 1 for "Tau4RD fibril polymorphism is imprinted during early aggregation"

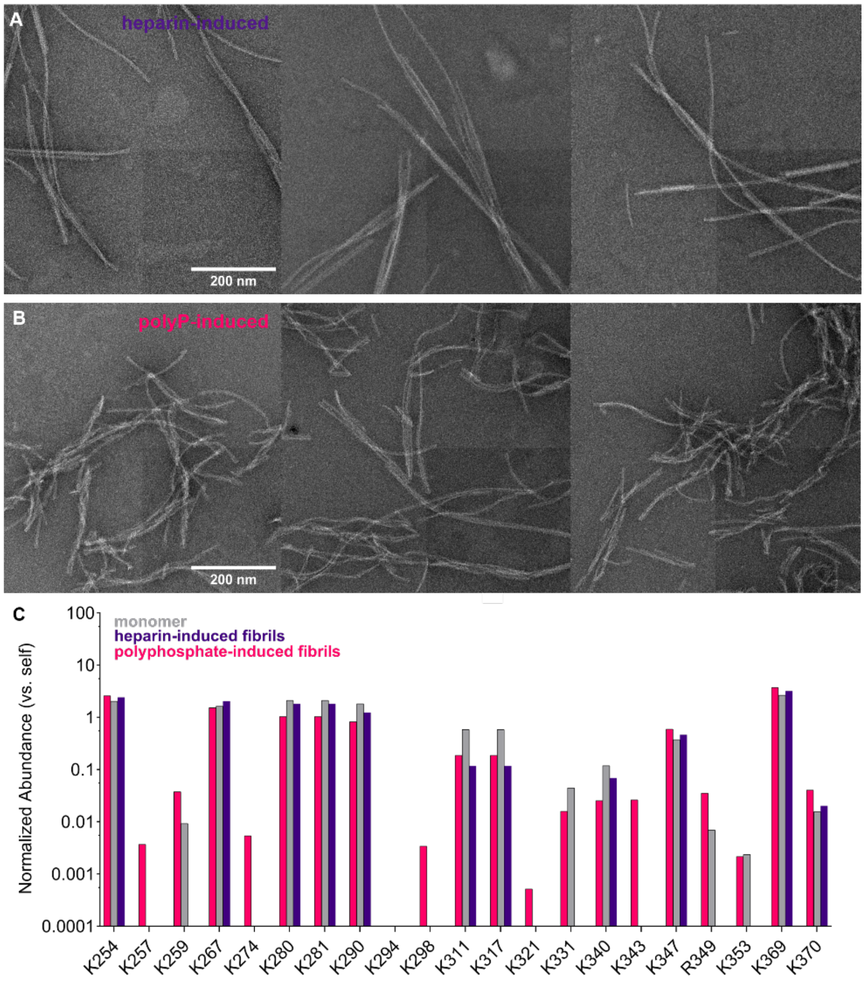


**SI Fig. 1: Differences in morphology between fibrils induced by heparin and polyphosphate.** A/B) Additional representative nsEM of heparin- and polyphosphate-induced tau4RD fibrils. Scale bar = 200 nm. Median filter; 4 px/6.4Å. C) Trypsin limited proteolysis of tau4RD monomer and heparin- or polyphosphate-induced fibrils reveals different cleavage fingerprints, providing further confirmation that heparin- and polyphosphate-generated fibrils are conformationally distinct.

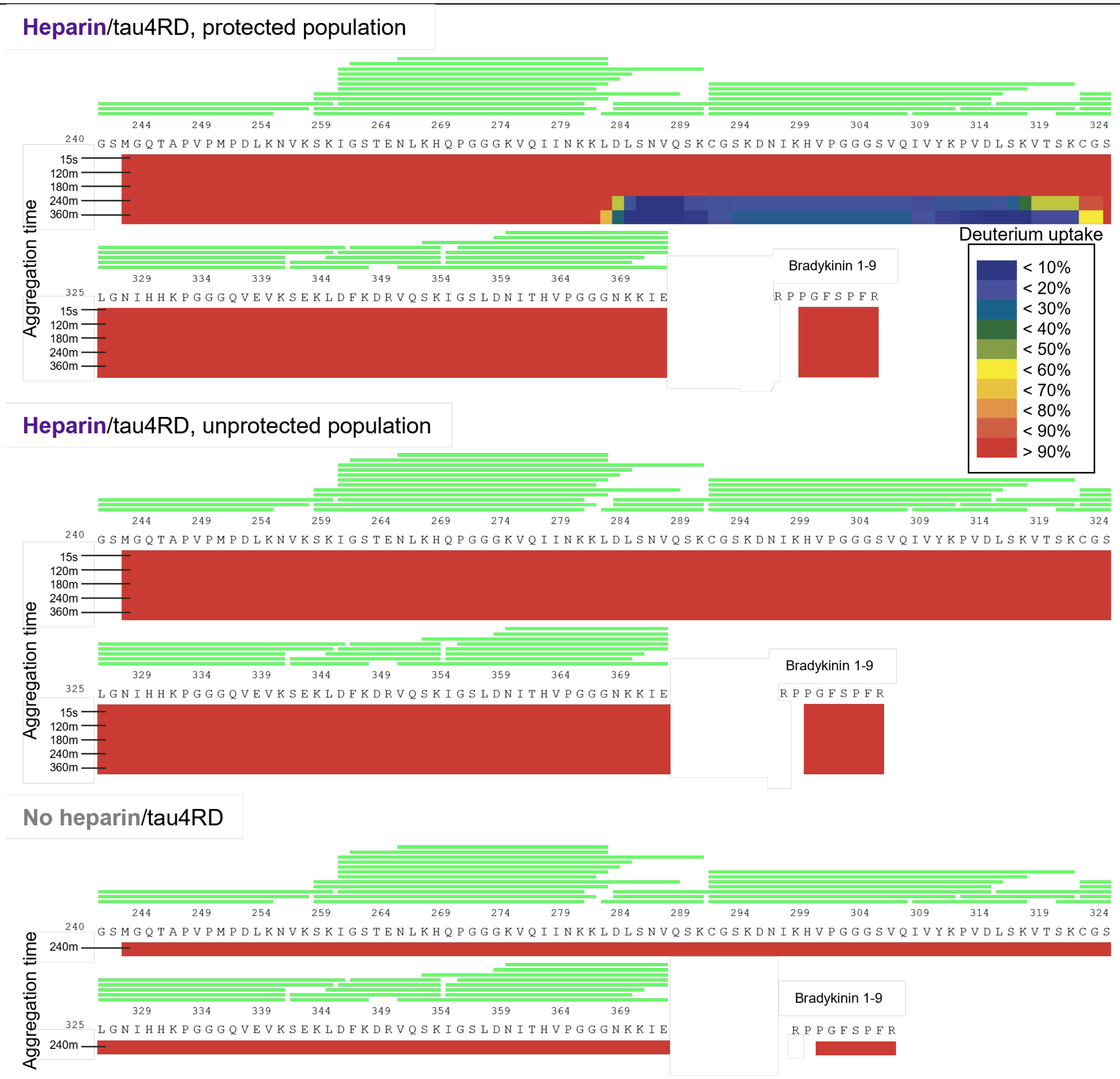


**SI Fig. 2: Deuterium uptake heatmap for heparin-induced tau4RD throughout aggregation.** Deuteration level per residue of tau4RD following a 3-sec 70% D2O pulse at various aggregation durations. This is a traditional visualization of the data displayed in Fig. 2A. Without inducer (no heparin condition), tau4RD monomer becomes completely deuterated with a 3-sec pulse. As aggregation proceeds, protection develops between residues 283-324. Note that the contribution of the protected and unprotected populations could not be completely deconvoluted in this analysis software (HD Examiner), so further fitting was performed on a subset of peptides with HXExpress v3.

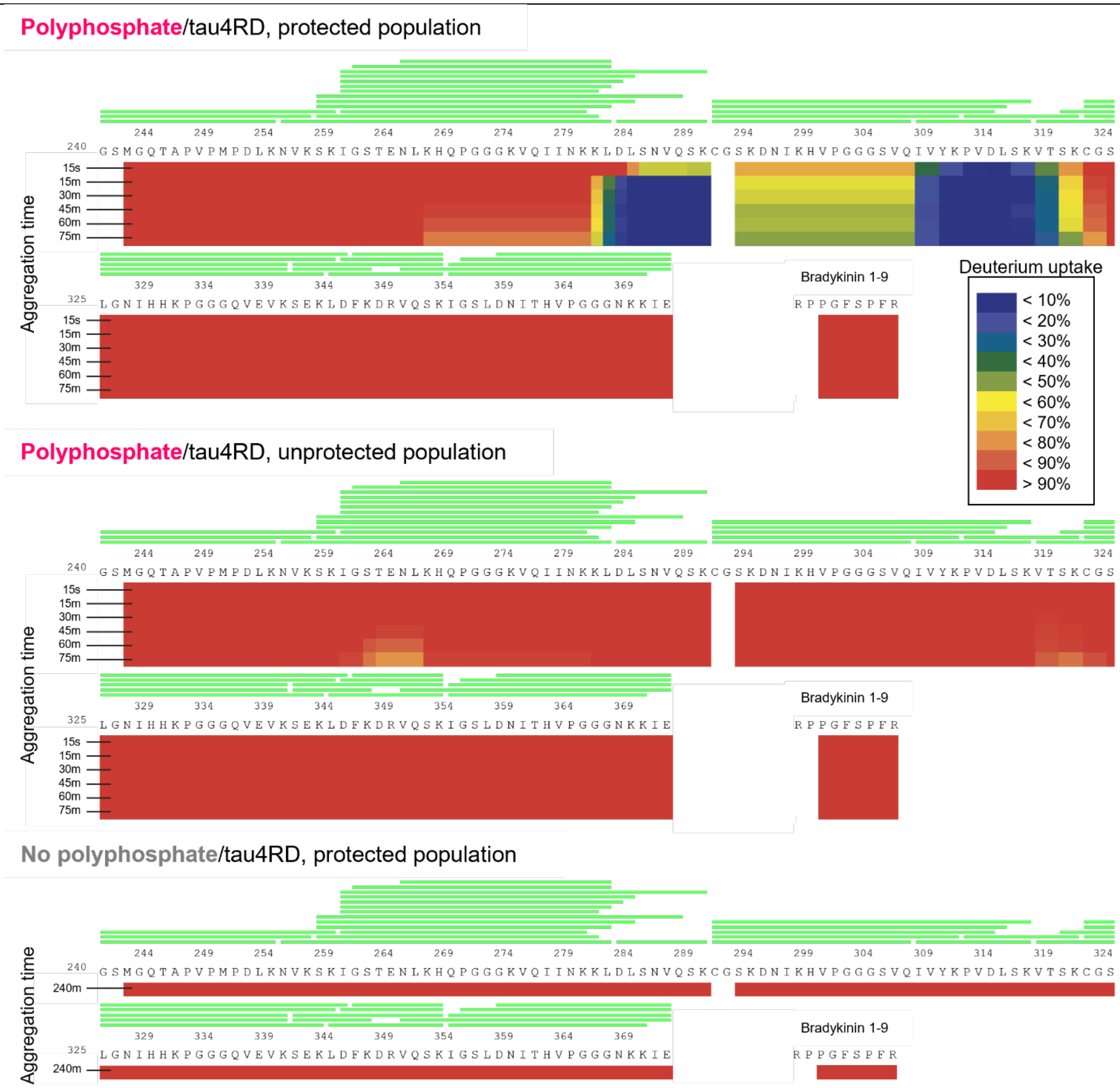


**SI Fig. 3: Deuterium uptake heatmap for polyphosphate-induced tau4RD throughout aggregation.** Deuteration level per residue of tau4RD following a 3-sec 70% D2O pulse at various aggregation durations. This is a traditional visualization of the data displayed in Fig. 2A. Without inducer (no polyphosphate condition), tau4RD monomer becomes completely deuterated with a 3-sec pulse. As aggregation proceeds, protection develops between residues 267-324. Note that the contribution of the protected and unprotected populations could not be completely deconvoluted in this analysis software (HD Examiner), so further fitting was performed on a subset of peptides with HXExpress v3.

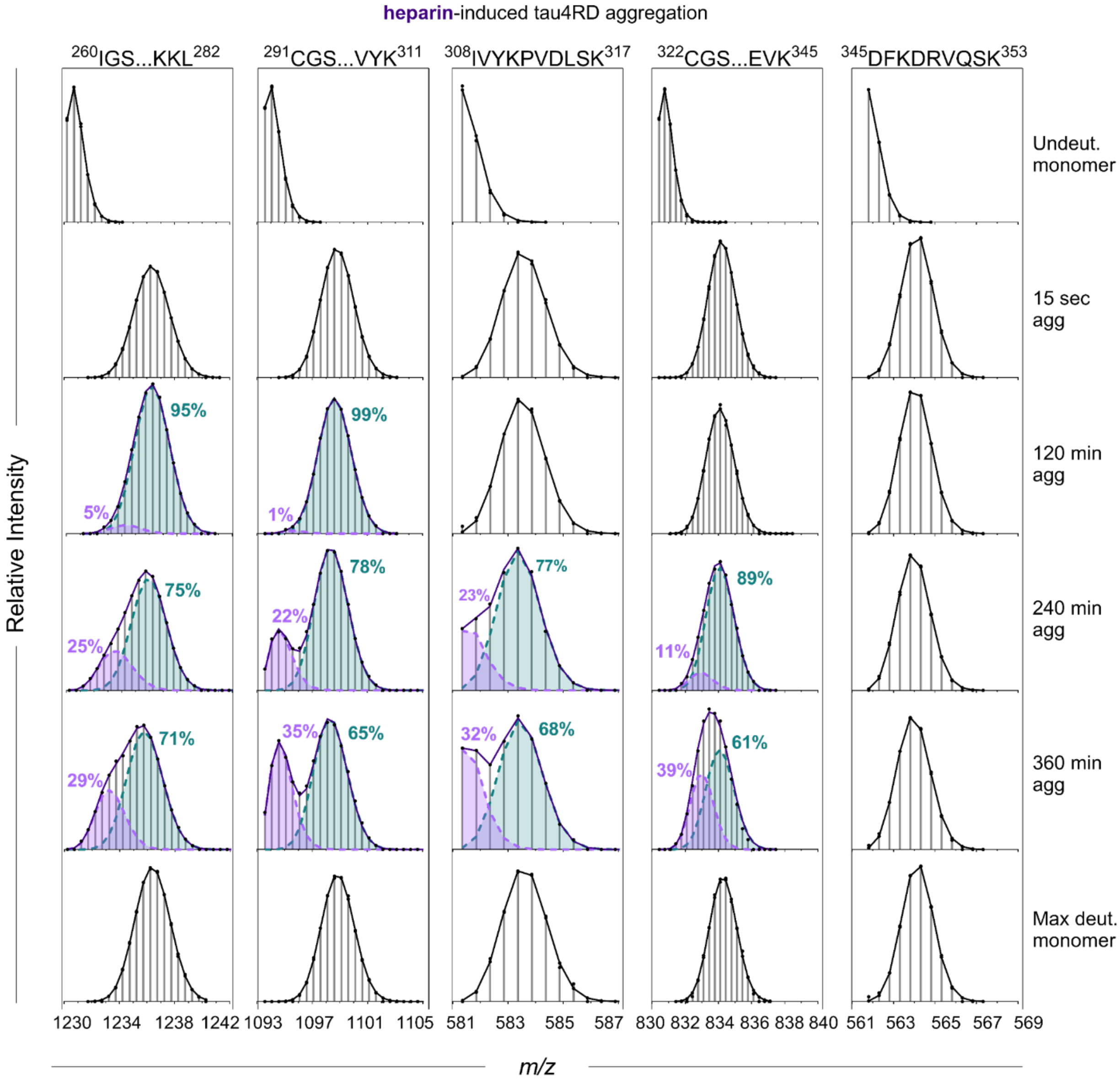


**SI Fig. 4: Mass envelopes for representative peptides over the course of heparin-induced tau4RD fibril formation.** Deuteration level per residue of tau4RD following a 3-sec 70% D2O pulse at various aggregation durations. This is a traditional visualization of the data displayed in Fig. 2A. Without inducer (no polyphosphate condition), tau4RD monomer becomes completely deuterated with a 3-sec pulse. As aggregation proceeds, protection develops between residues 267-324. Note that the contribution of the protected and unprotected populations could not be completely deconvoluted in this analysis software (HD Examiner), so further fitting was performed on a subset of peptides with HXExpress v3.

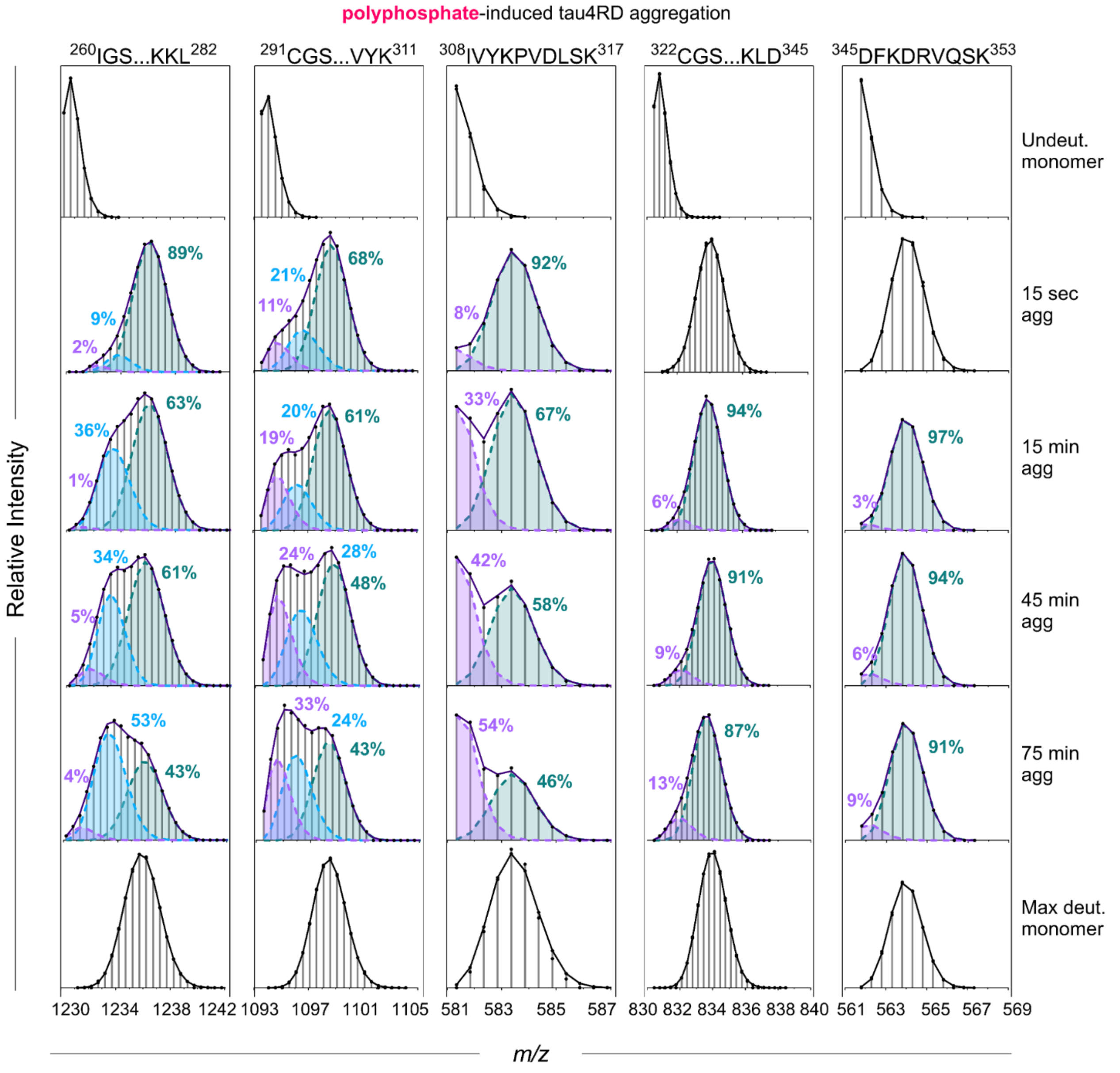


**SI Fig. 5: Mass envelopes for representative peptides over the course of polyphosphate-induced tau4RD fibril formation.** Deuteration level per residue of tau4RD following a 3-sec 70% D2O pulse at various aggregation durations. This is a traditional visualization of the data displayed in Fig. 2A. Without inducer (no polyphosphate condition), tau4RD monomer becomes completely deuterated with a 3-sec pulse. As aggregation proceeds, protection develops between residues 267-324. Note that the contribution of the protected and unprotected populations could not be completely deconvoluted in this analysis software (HD Examiner), so further fitting was performed on a subset of peptides with HXExpress v3.

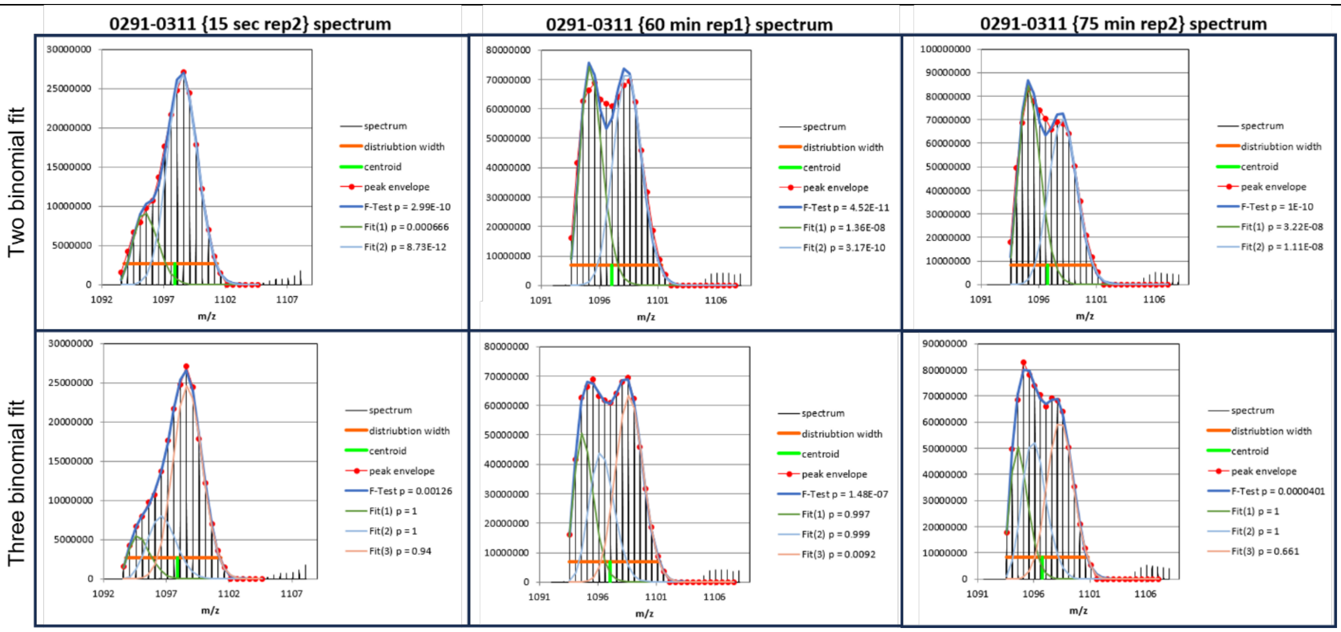


**SI Fig. 6: Statistical support for multimodal (three binomial) fits for polyphosphate-induced tau4RD peptide $^{291}$CGS…VYK$^{311}$.** Mass spectra of peptide $^{291}$CGS…VYK$^{311}$ at t = 15 sec, 60 min, 75 min after aggregation initiation. Mass envelopes have been fit to two binomials in the top row and three binomials in the bottom row. While the subpopulations cannot be accurately deconvoluted with a three-binomial fit, a two-binomial fit does not sufficiently describe the data (e.g., mass envelope not accounted for in the two-binomial fit of 60 min rep1). Note that the F-test p-value remains statistically significant after three binomial fits, even with the substantial penalty applied for adding an additional degree of freedom. Fitting and plots from HXExpress v3.

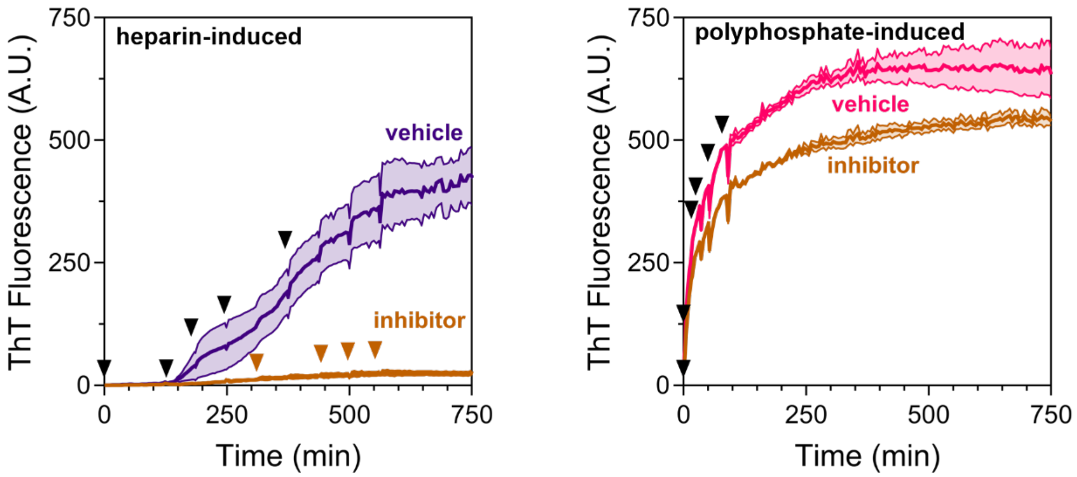


**SI Fig. 7: Tryptanthrin inhibits heparin- and polyphosphate-induced tau4RD aggregation.** ThT traces for heparin- (left) and polyphosphate-induced (right) tau4RD aggregation in the presence of inhibitor (1:2) or vehicle control. Aggregation inhibitor increases the lag time and strongly decreases the plateau height of heparin-induced tau4RD aggregation. Addition of inhibitor to polyphosphate-induced tau4RD aggregation results in no change to the lag time but results in a moderate decrease of the plateau height.

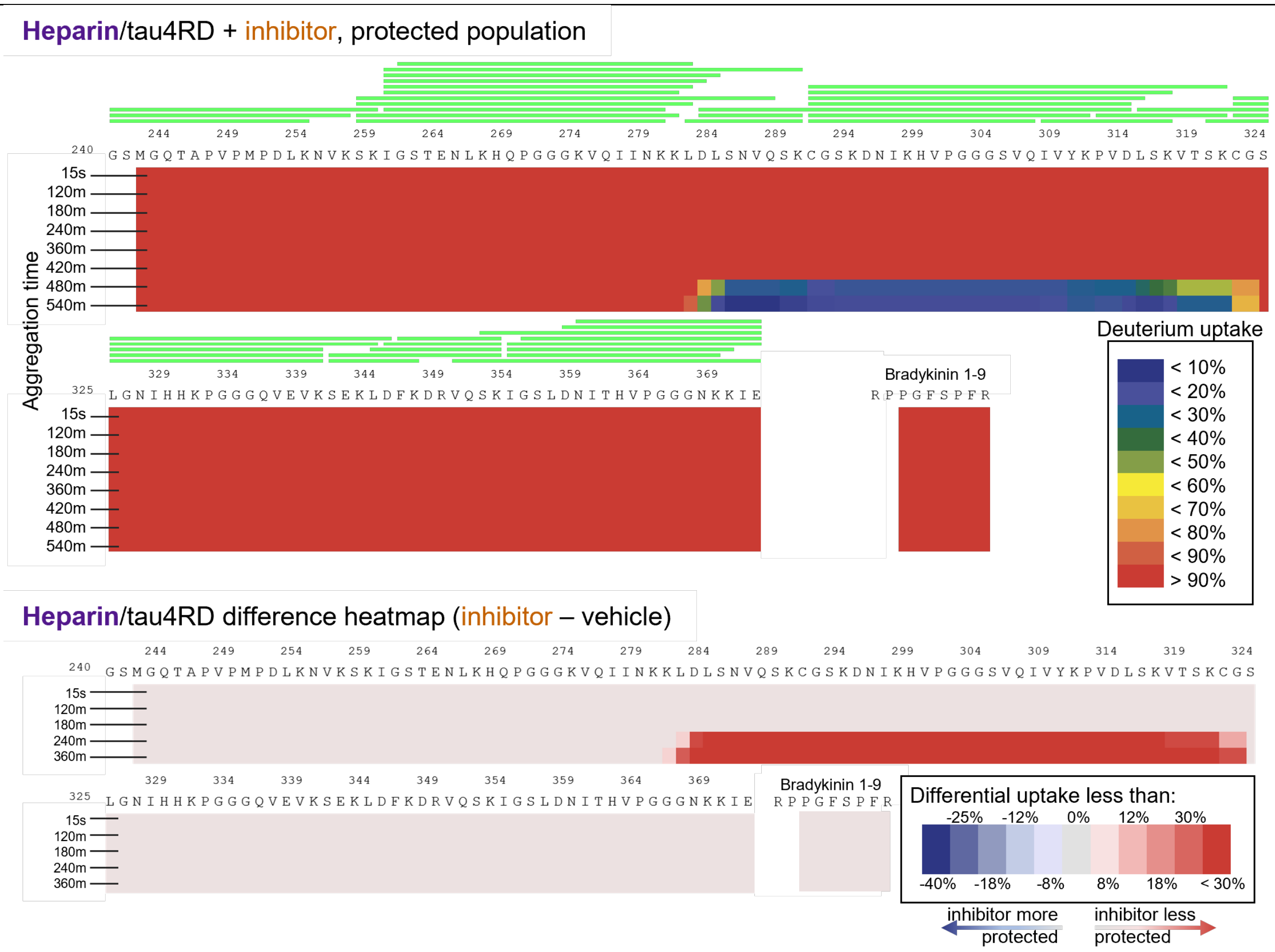


**SI Fig. 8: Deuterium uptake heatmap for heparin-induced tau4RD incubated with tryptanthrin (inhibitor).** Top) Deuteration level per residue of tau4RD following a 3-sec 70% D2O pulse at various aggregation durations. This is a traditional visualization of the data displayed in Fig. 4A. As aggregation proceeds, protection develops between residues 267-324. Note that the contribution of the protected and unprotected populations could not be completely deconvoluted in this analysis software (HD Examiner), so further fitting was performed on a subset of peptides with HXExpress v3. Bimodal populations were detected at 15-sec (thought to be caused by inhibitor binding) after HXExpress deconvolution. See SI Figs. 12 and 14. Bottom) Difference heatmap showing that the inhibited condition is less protected (less aggregated) at 240 m and 360 m.

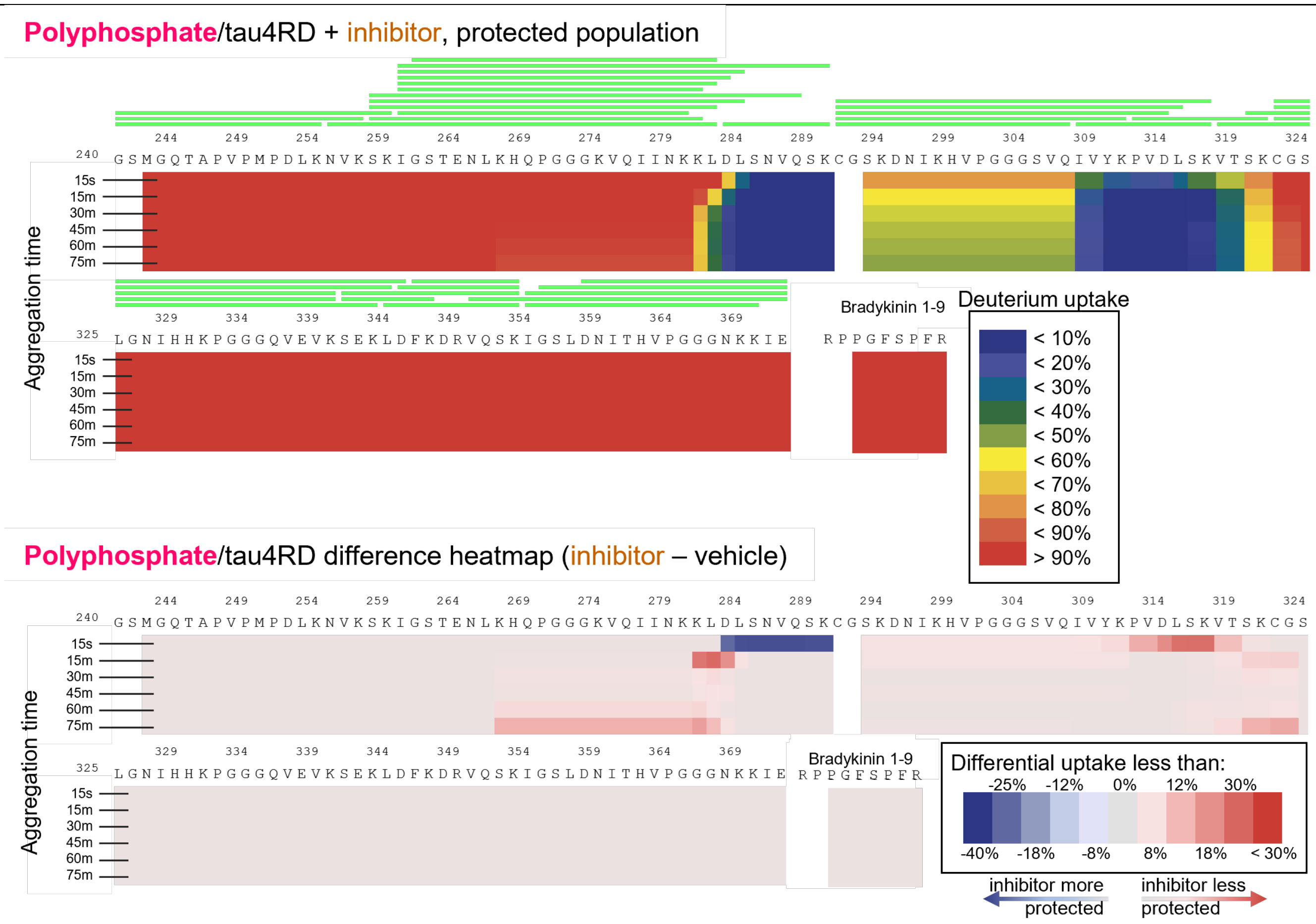


**SI Fig. 9: Deuterium uptake heatmap for polyphosphate-induced tau4RD incubated with inhibitor.** Top) Deuteration level per residue of tau4RD following a 3-sec 70% D2O pulse at various aggregation durations. This is a traditional visualization of the data displayed in Fig. 5A. As aggregation proceeds, protection develops between residues 267-324. Note that the contribution of the protected and unprotected populations could not be completely deconvoluted in this analysis software (HD Examiner), so further fitting was performed on a subset of peptides with HXExpress v3. Bottom) Difference heat map showing that tau4RD incubated with inhibitor is generally slightly less protected (less aggregated) than the vehicle control. Note the exception between residues 284-290, which exhibit substantial protection in the presence of inhibitor at the 15-sec time point. We attribute this protection to inhibitor binding.

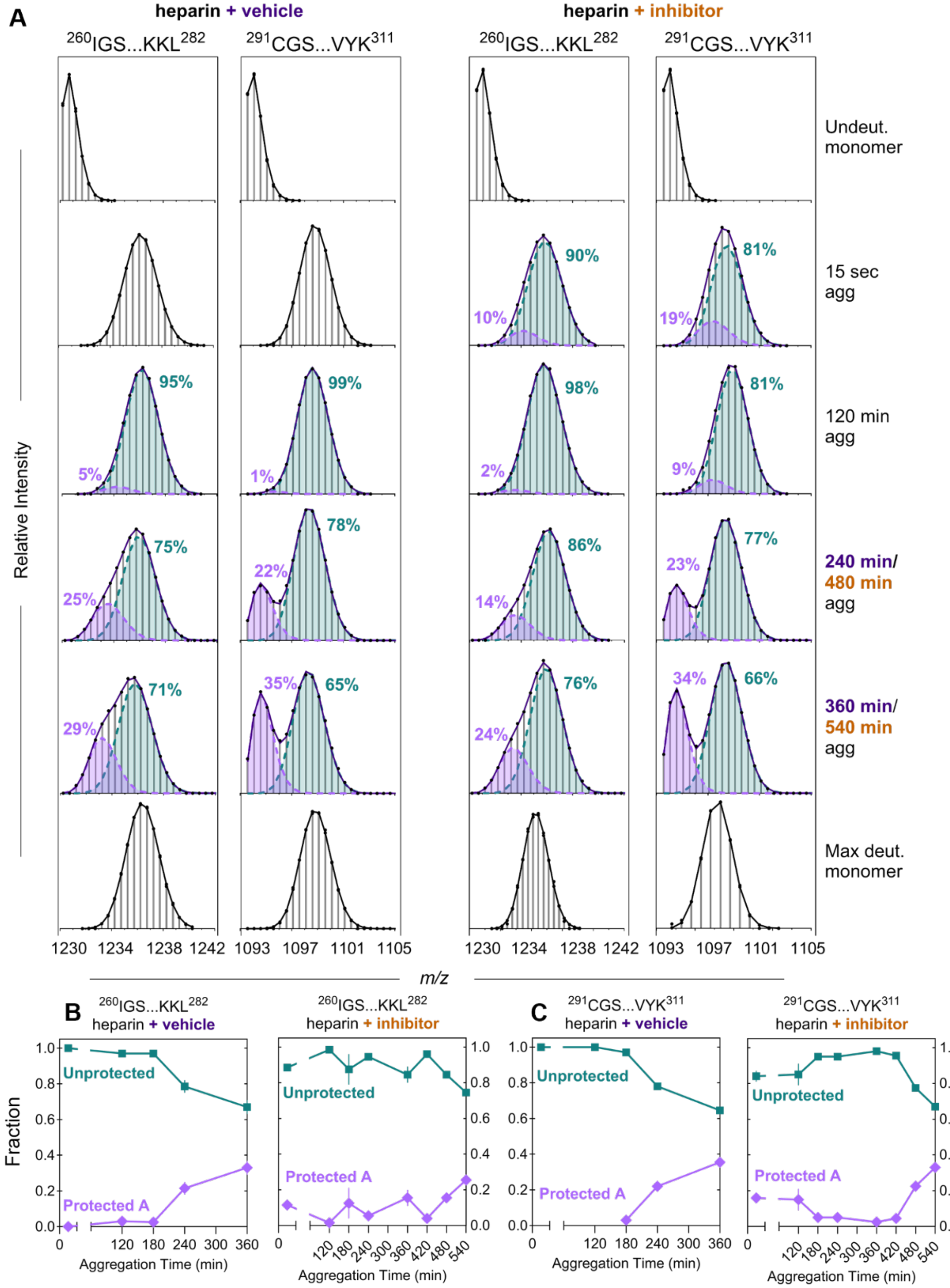


**SI Fig. 10: Multimodal behavior observed in the presence of an inhibitor during heparin-induced tau4RD aggregation.** A) Data for two peptides in the absence (vehicle) or presence of tryptanthrin (inhibitor). Although the addition of tryptanthrin substantially delays and reduces heparin-induced tau4RD aggregation, it does not result in changes to the fibril core. F-test p-value is less than 0.05 for all mass envelopes fit with multiple binomials. B/C) Population fraction for the aggregated (protected) and monomer-like (unprotected) populations for peptides I260-L282 and C291-K311.

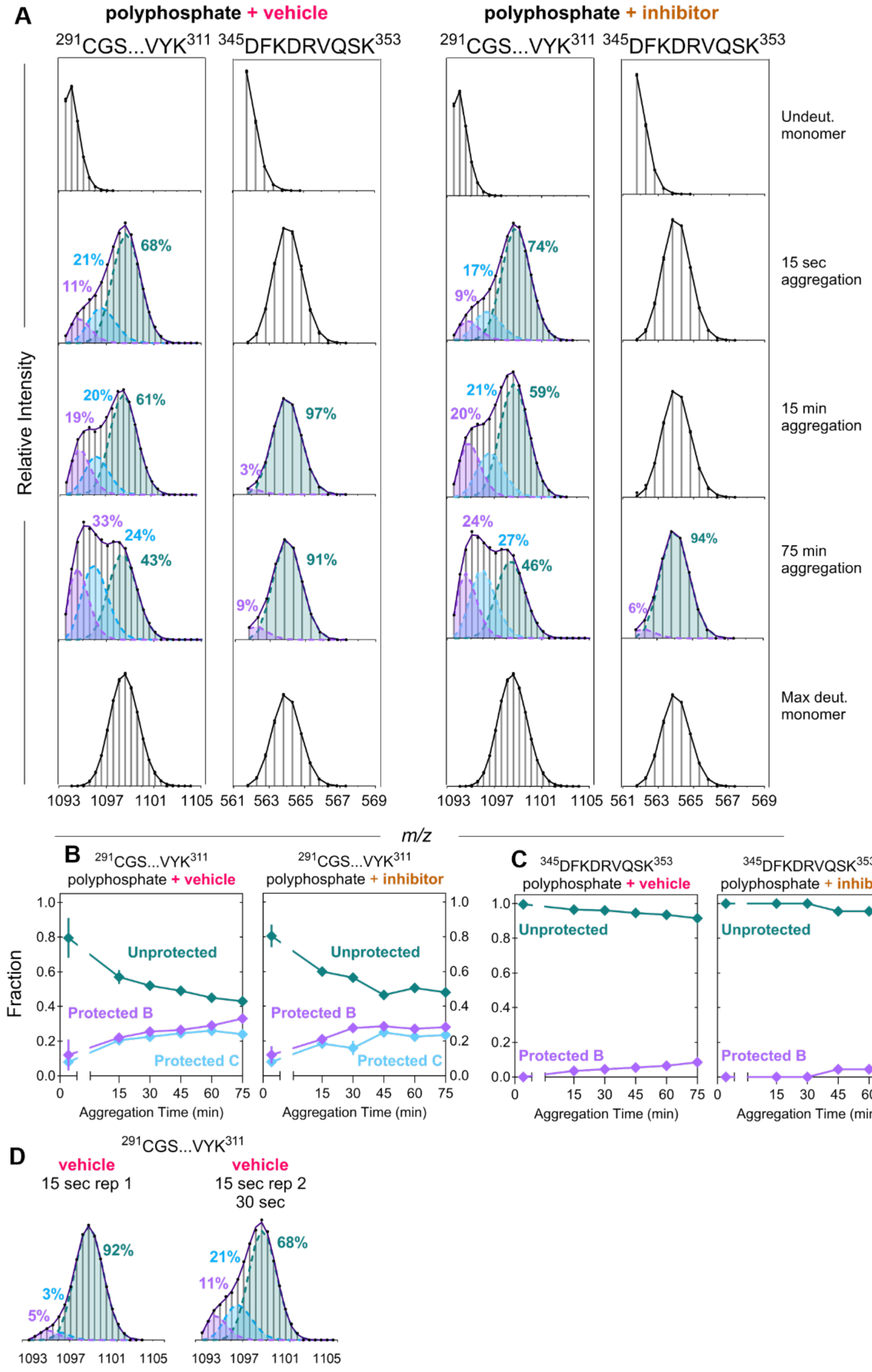


**SI Fig. 11: Multimodal behavior observed in the presence of an inhibitor during polyphosphate-induced tau4RD aggregation.** A) Data for two peptides in the absence (vehicle) or presence of tryptanthrin (inhibitor). Although the addition of tryptanthrin mildly reduces polyphosphate-induced tau4RD aggregation, it does not result in changes to the fibril core. F-test p-value is less than 0.05 for all mass envelopes fit with multiple binomials. B/C) Population fraction for the aggregated (protected) and monomer-like (unprotected) populations for peptides C291-K311 and D345-K353. D) Duplicate sampling at 15 sec results in substantially different subpopulation profiles due to the speed at which polyphosphate-induced aggregation progresses.

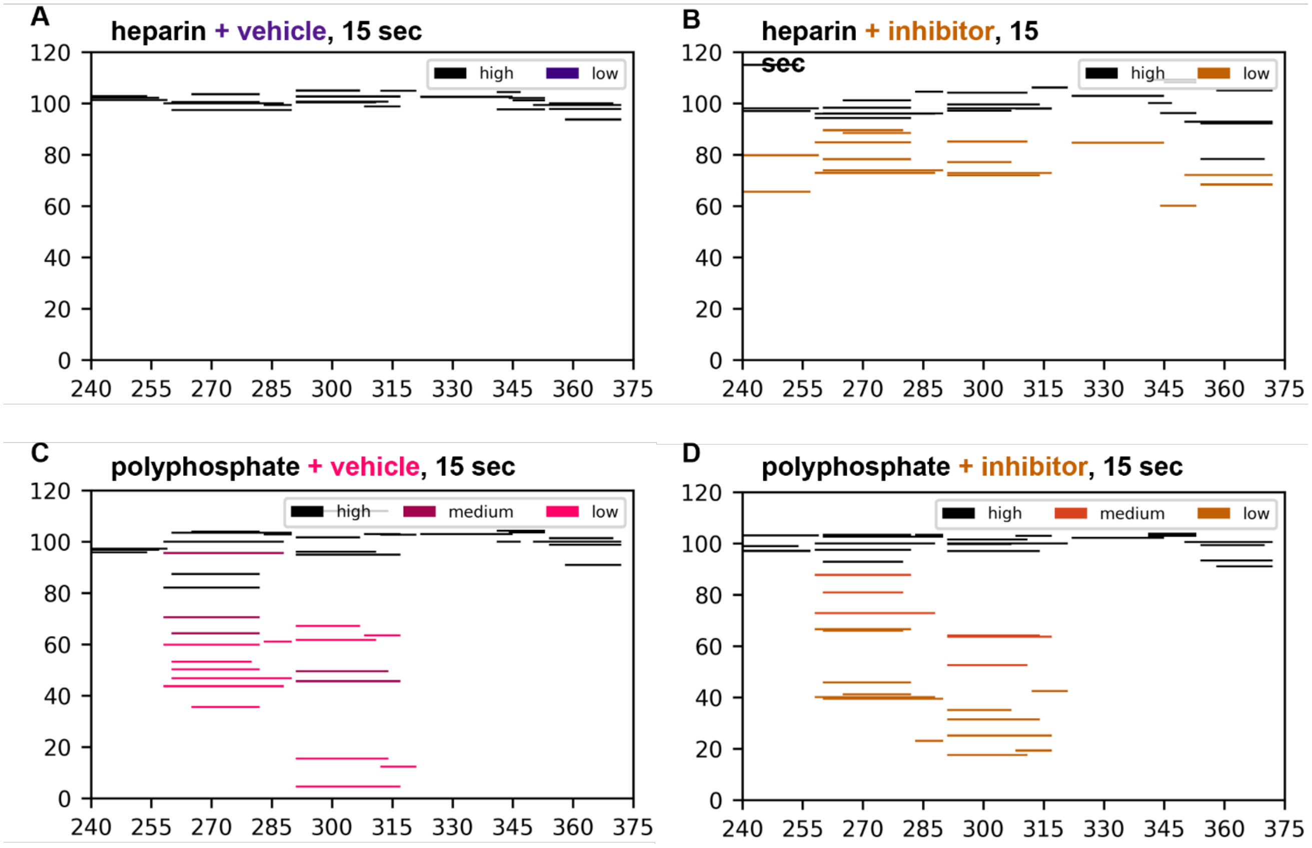


**SI Fig. 12: HDX Woods plots for heparin- and polyphosphate-induced tau4RD incubated with inhibitor for 15-sec.** A) Woods plot revealing only a high-exchanging (monomer-like) population when heparin-induced tau4RD is incubated with vehicle. B) In contrast, a Woods plot of heparin-induced tau4RD incubated with inhibitor shows both a high-exchanging (monomer-like) and low-exchanging (protected) population. The pattern of protection caused by the inhibitor does not match the pattern of protection caused by the heparin fibril core, suggesting that the protection observed at 15-sec is due to structural changes associated with the inhibitor binding tau4RD. C) Woods plot revealing both morph B and morph C are present 15-sec after the beginning of polyphosphate-induced tau4RD aggregation. D) The addition of inhibitor to polyphosphate-induced tau4RD aggregation does not change the pattern of protection.

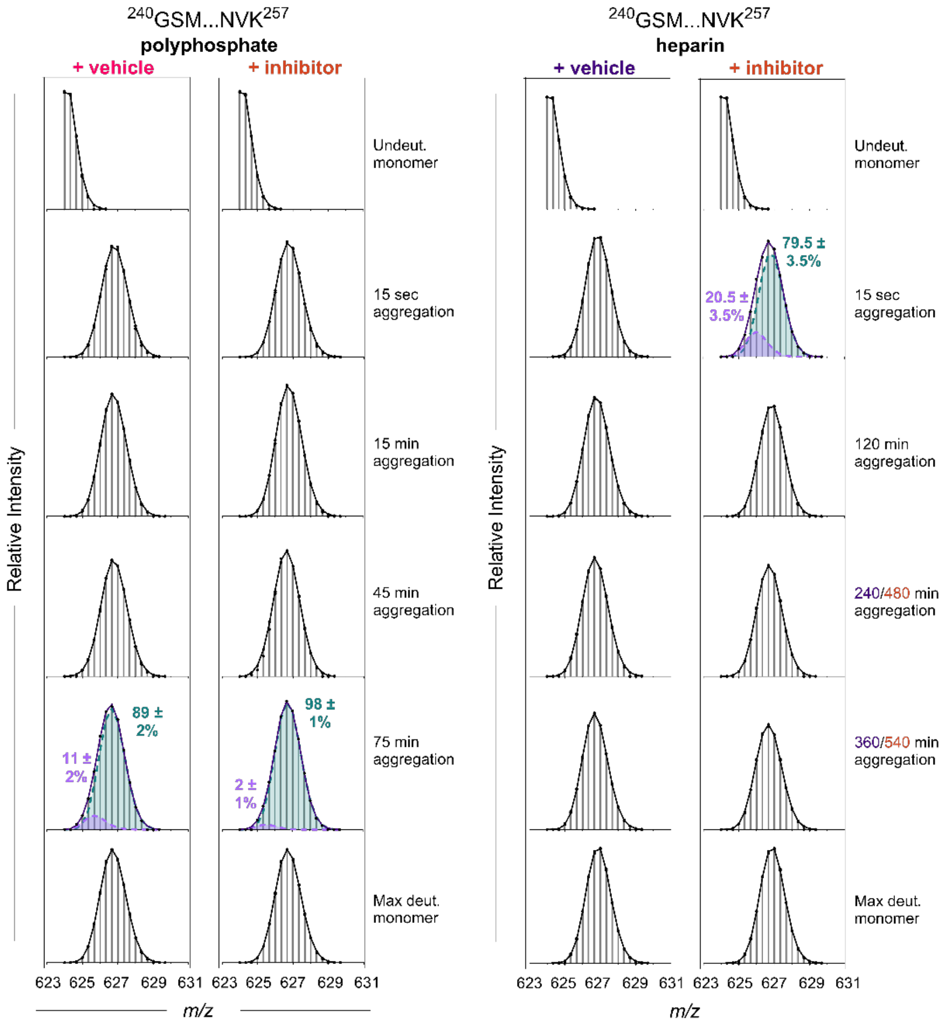


**SI Fig. 13: Addition of inhibitor does not change the low abundance aggregated subpopulation observed in peptide 240-257 under polyphosphate-initiated aggregation.** A) Data for one peptide in the absence (vehicle) or presence of tryptanthrin (inhibitor) when tau4RD aggregation is induced with polyphosphate. Note that the low-abundance population is not detected until 75 min in either case. This likely represents fibril maturation. F-test p-value is less than 0.05 for all mass envelopes fit with multiple binomials. B) Peptide 240-257 when incubated in the absence (vehicle) or presence (inhibitor) of tryptanthrin. Note that there appears to be a low abundance protected population at 15 sec in the presence of inhibitor; this could be due to inhibitor binding and causing rearrangements in the tau4RD backbone. Note that there is no evidence of an aggregated subpopulation at the final time point (360 min for vehicle, 540 min for inhibitor). Subpopulation abundance values are the average (mean) ± difference between duplicate samples.

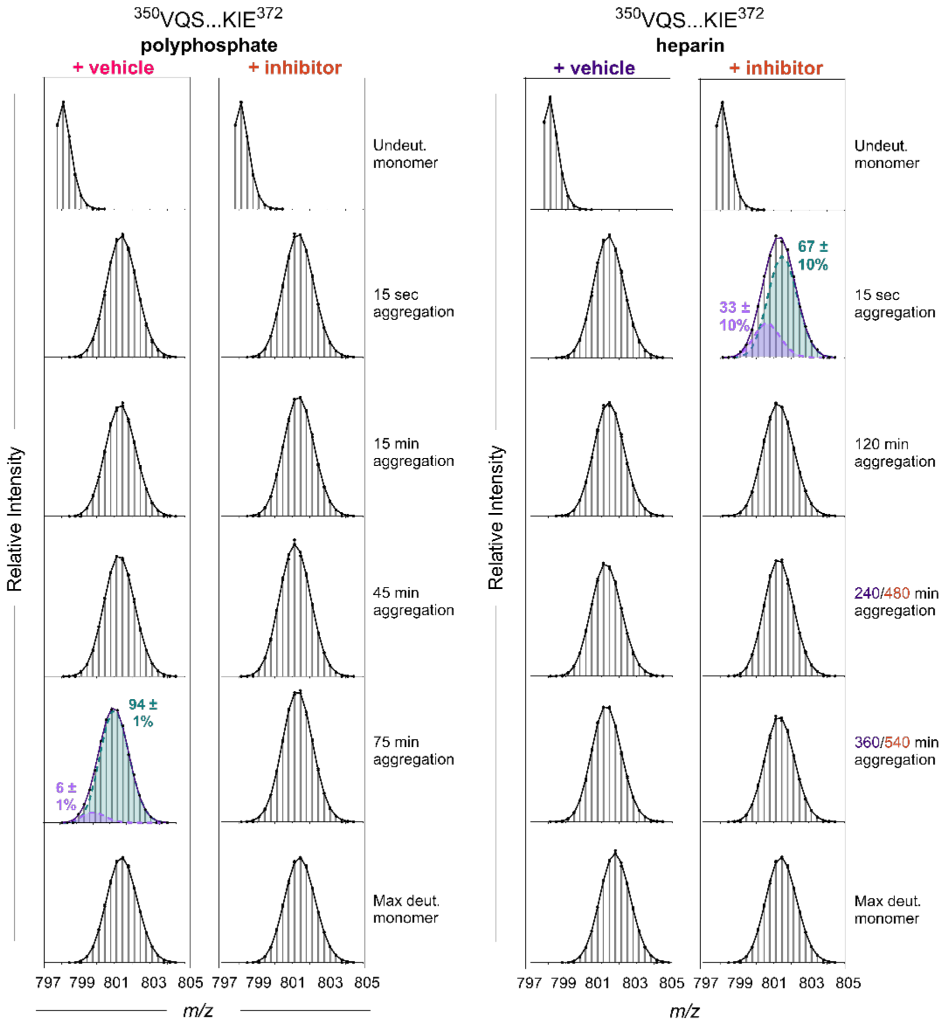


**SI Fig. 14: Inhibitor reduces the low abundance aggregated subpopulation observed during polyphosphate-induced tau4RD aggregation at peptide 350-372.** A) Spectra for peptide 350-372 with vehicle or inhibitor when tau4RD aggregation is induced with polyphosphate. Note that the low-abundance population is not detected until 75 min with vehicle. This likely represents fibril maturation. There is not evidence of an aggregated subpopulation at 75 min when tau4RD is incubated with tryptanthrin. This is likely due to a reduction in overall aggregation and not due to a conformational change caused by tryptanthrin. F-test p-value is less than 0.05 for all mass envelopes fit with multiple binomials. B) Peptide 350-372 incubated with vehicle or inhibitor and aggregation is induced with heparin. Note that there appears to be a low abundance protected population at 15 sec in the presence of inhibitor; this could be due to inhibitor binding and causing rearrangements in the tau4RD backbone. Note that there is no evidence of an aggregated subpopulation at the final time point (360 min for vehicle, 540 min for inhibitor). Subpopulation abundance values are the average (mean) ± difference between duplicate samples.
